# Cell-type-specific firing patterns in a V1 cortical column model depend on feedforward and feedback-driven states

**DOI:** 10.1101/2024.04.02.587673

**Authors:** Giulia Moreni, Rares A. Dorcioman, Cyriel M. A. Pennartz, Jorge F. Mejias

## Abstract

Stimulation of specific cell groups under different network regimes (e.g., spontaneous activity or sensory-evoked activity) can provide insights into the neural dynamics of cortical columns. While these protocols are challenging to perform experimentally, modelling can serve as a powerful tool for such explorations. Using detailed electrophysiological and anatomical data from mouse V1, we built a novel spiking network model of a cortical column, which incorporates pyramidal cells and three distinct interneuron types (PV, SST, and VIP cells, specified per lamina), as well as the dynamic and voltage-dependent properties of AMPA, GABA, and NMDA receptors. We first demonstrate that thalamocortical feedforward (FF) and feedback (FB) stimuli arriving in the column have opposite effects, leading to net columnar excitation and inhibition respectively and revealing translaminar gain control via full-column inhibition by layer 6. We then perturb one cell group (i.e. a cell type in a specific layer) at a time and observe the effects on other cell groups under distinct network states: spontaneous, feedforward-driven, feedback-driven, and a combination of feedforward and feedback. Our findings reveal that when a given group is perturbed, the columnar response varies significantly based on its state, with strong sensory feedforward input decreasing columnar sensitivity to all perturbations and feedback input serving as modulator of intra columnar interactions. Given that activity changes within specific neuronal populations are difficult to predict a priori in experiments, our model may constitute a useful tool to predict outcomes of perturbations and assist in experimental design.

**Author Summary:** In this study, we explored how stimulating specific groups of neurons under different conditions (spontaneous or stimulus-evoked states) can help us understand the dynamics of visual cortical columns in mouse cortex. We developed a data-constrained model of a cortical column that includes several cell types (excitatory neurons as well as multiple inhibitory neuron types) and synaptic receptors. Our findings reveal that signals coming from higher brain areas (feedback input) and sensory input (feedforward input) have different impacts on the column: sensory inputs generally increase activity within the column, while feedback tends to decrease it. We then considered input targeting one cell type at a time to see how it affects the internal dynamics. Our results show that neural responses to selected input vary depending on whether the cortical column is in a resting state or being stimulated by feedforward input or feedback input. This model is a useful tool for predicting the outcomes of these cell interactions, which can help in planning real-world experiments and understanding how our brain processes information.

## Introduction

The intricate architecture of the cerebral cortex, with its diverse neuronal populations and complex connectivity patterns, plays a pivotal role in processing sensory information and orchestrating cognitive functions [1,2]. Central to this architecture is the column-like circuitry, a functional unit that has been the subject of extensive research over the years [3–5]. Within these columns, different neuronal populations interact in a dynamic manner [6], giving rise to a rich repertoire of activity patterns that can be modulated by both internal and external factors [7]. One of the intriguing aspects of cortical columns is the interplay between excitatory and inhibitory neuronal populations. This balance, often referred to as E/I balance, is crucial for maintaining stable network dynamics and ensuring efficient information processing [8–13]. Disruptions in this balance can lead to various neurological and psychiatric disorders (such as schizophrenia and autism) underscoring its importance [14,15]. In recent years, there has been a growing interest in understanding how different types of input, such as cortical and thalamic feedforward (FF) and feedback (FB) signals, modulate the activity within cortical columns [16–19]. These inputs can have profound effects on the firing rates of different neuronal populations, leading to diverse patterns of network activity [16,19]. However, the precise mechanisms through which these inputs shape cortical dynamics remain a topic of active investigation [20,21].

In this study, we delve into the behaviour of a cortical column model under different conditions. We explore how spontaneous activity, feedforward input and feedback input influence the activity of different neuronal populations within the cortical column. This was done by building a novel biologically detailed computational model of a cortical column of mouse primary visual cortex V1. This model incorporates dynamic properties of multiple postsynaptic receptors (AMPA, NMDA, and GABA-A receptors), and is tightly constrained by state-of-the-art cortical connectivity data on mouse V1 [22,23] which include cell densities and laminar-specific connectivity patterns across four different cell types (pyramidal neurons and PV, SST and VIP interneurons) and five laminar modules (layers 1, 2/3, 4, 5 and 6). Our simulations show that feedforward and feedback stimuli have opposing impacts on columnar activity, leading to net columnar excitation and inhibition, respectively, in agreement with experimental and other modelling work [19,24–26]. Furthermore, our model reveals a translaminar inhibitory effect mediated by layer 6 activity, in line with a role in columnar gain control role as suggested by experiments [27,28]. However, our model reveals that such inhibition may well be the consequence of complex cortical interactions and not only of the recruitment of local inhibitory cells as previously suggested. We then stimulated one cell group at a time in distinct states (spontaneous, sensory feedforward-driven, feedback-driven, and a state driven by a combination of feedforward and feedback stimuli) and observed the effects on the other populations. These pertubational experiments reveal multiple group-to-group dependencies which vary across the different states considered, with strong feedforward input decreasing overall columnar sensitivity and feedback input modulating interactions linked lateral, inter-columnar communication. In addition, our findings proved the distinctive roles of inhibitory neurons in network dynamics. For instance, our model predicts that in an experimental setting with strong top-down modulation of V1—such as input to excitatory neurons in layer 5—providing stimuli to different inhibitory neurons in layer 6 will induce contrasting responses in the E2/3 neurons, depending on the specific inhibitory subgroup targeted. This model illuminate the complex interplay between different types of inputs and their impact on cortical dynamics and it can be used as a predictive tool to guide and design experiments. Moreover, our results reveal that when the same group is stimulated, the network’s response can vary significantly based on its initial state.

## Results

We begin by describing our cortical column model, sketched in Fig. 1A. We consider a network of 5,000 spiking neurons which are distributed across five laminar modules, four of which (layers 2/3, 4, 5 and 6) contain pyramidal neurons as well as PV, SST and VIP cells, and one of which (layer 1) contains only VIP cells. For laminar modules with multiple cell types, about 85% of the cells are pyramidal neurons and the remaining 15% are inhibitory interneurons –with the precise proportion of each inhibitory cell type in each layer given by anatomical data^23^ and depicted in Fig. 1A as the relative size of the respective inhibitory population.

**Figure 1:**
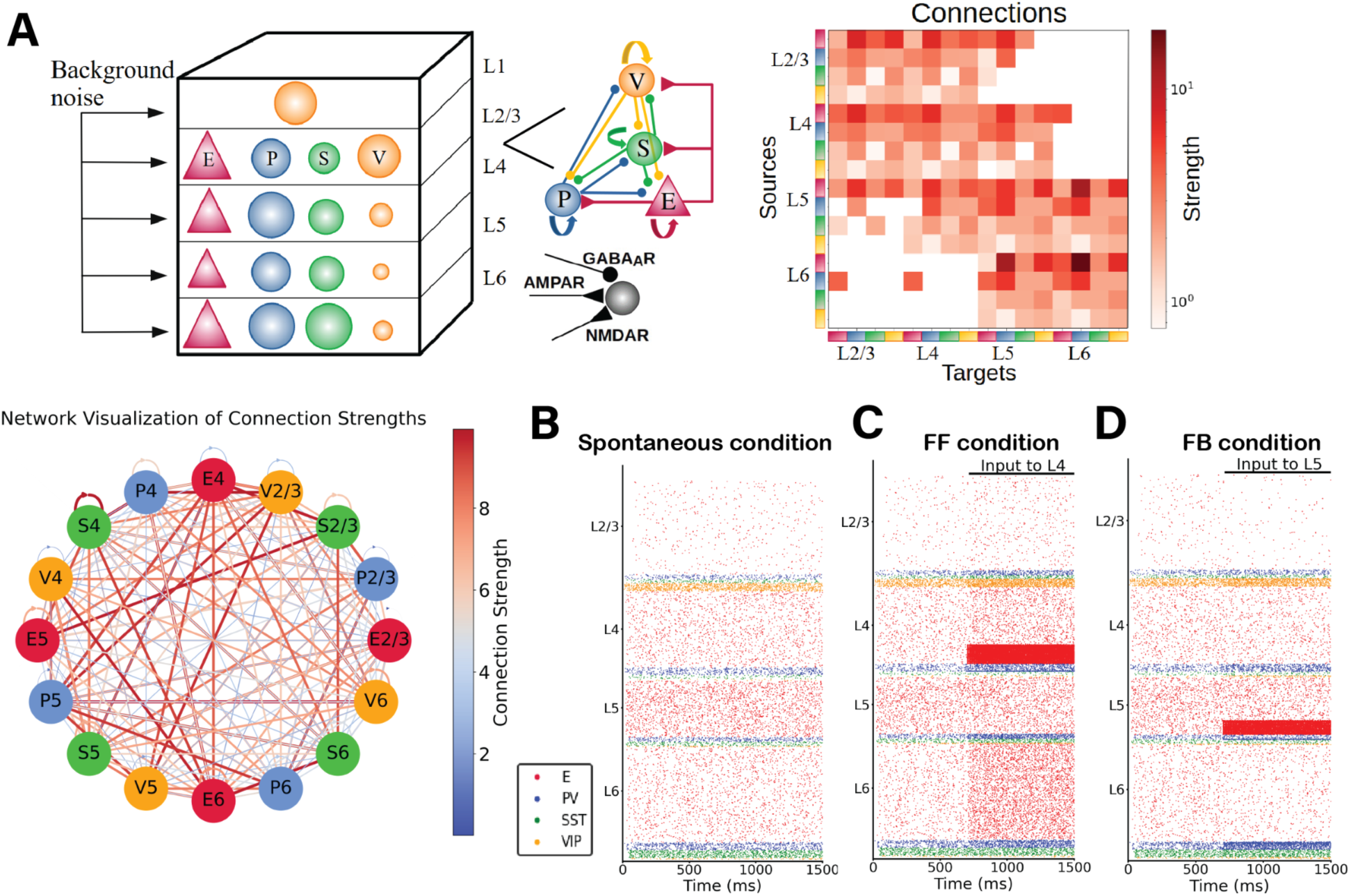
(A) Sketch of the cortical column model. In layers 2/3, 4, 5, and 6 an excitatory population E (red triangles) and 3 types of inhibitory population (PV, SST, VIP cells as blue, green, orange circles: P, S and V, respectively) are present. In layer 1 only VIP cells are present. The size of the circles in the top-left panel represents the relative size of the inhibitory populations. Connections between groups are not shown in the top left diagram; the zoomed-in schematic to the right shows inter-population connectivity and postsynaptic receptors (AMPA, GABA_A_, NMDA) involved. The connectivity matrix is shown at the right and visualized in the network-like sketch. (B)-(D) Network in three different states. (B) Spontaneous (resting) condition. Simulated firing rates match experimental firing rates [21]. (C) Feedforward-driven condition (i.e., with excitatory input to L4 cells). At 700ms an constant input of 150 pA is injected to 25% of L4 pyramidal cells and 5% of PV cells. Excitatory neurons in all layers increase their activity. (D) One feedback-driven condition (i.e., with excitatory input to L5 pyramidal cells). At 700ms an input of 150pA is injected to 25% of pyramidal cells and 5% of PV cells in L5, and excitatory neurons in all other layers decrease their activity while interneurons in layer 6 increase their firing.

### Spontaneous cell type and layer-specific activity

To match the spontaneous firing rates of all cell types observed in vivo, we adjusted the global scaling value for the entire connectivity and the cell-specific background inputs to the column, similar to previous work [29,30]. The resulting simulated spontaneous, feedforward-evoked and feedback-evoked spiking activity in the cortical column is displayed as a raster plot in Fig. 1B. In the spontaneous state, for all cell types in our model, asynchronous irregular activity patterns were obtained, with firing rate levels matching quite closely those observed in vivo (Fig. 2A). The activity varied substantially across layers and cell types. Across all layers, pyramidal neurons exhibited the lowest firing rates in their laminar module, with mean firing rates around 2 Hz for layer 5 and below or close to 1 Hz for other layers, in agreement with experimental data [23]. In all layers, firing rates of inhibitory cells exceeded those of excitatory cells, except for VIP cells in layer 4. This sets the columnar model in an inhibition-dominated regime with a basal pattern of asynchronous firing [31].

**Figure 2:**
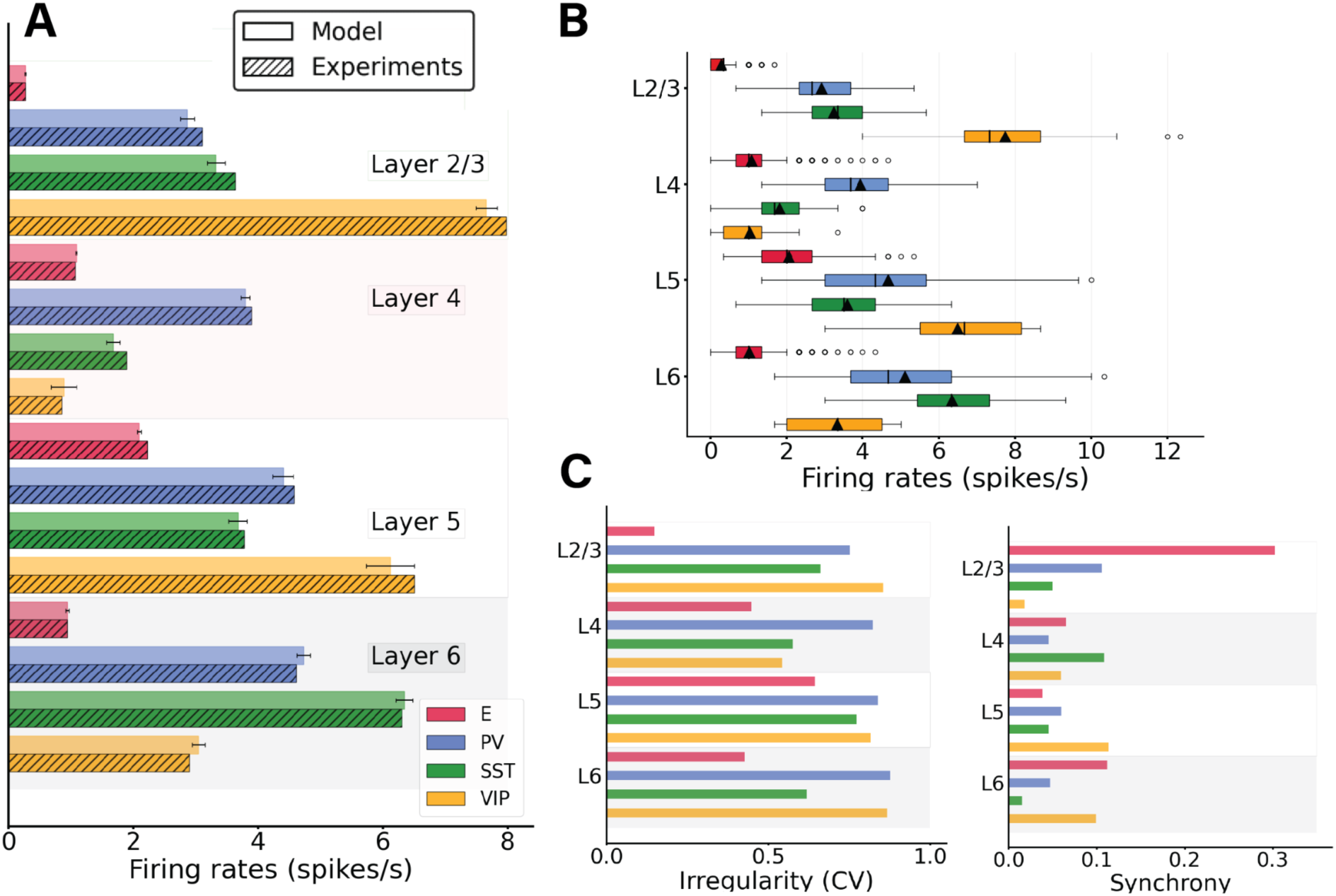
Spontaneous cell-type specific activity in the columnar model. (A) Mean firing rates for each model population (full bars, standard deviation computed over 10 network realisations or initializations) vs experiment (dashed bars). (B) Boxplot of single-unit firing rates in the model. Circles show outliers, black triangles indicate the mean firing rate of the population, and black vertical lines in each box indicate the median. (C) Left: Irregularity of single-unit spike trains quantified by the coefficient of variation of the inter-spike intervals. Right: Synchrony of multi-neuron spiking activity quantified by membrane potential traces.

In addition to the heterogeneity in firing rates found between layers and cell types, single-neuron firing rates within the same population also displayed substantial variability (Fig. 2B). For example, some pyramidal neurons in layer 2/3 fired at 2 Hz, while the majority of them were rather quiescent, emitting less than one spike per second. This is in agreement with previous findings [30]. Overall, single-unit activity was quite irregular, with the mean of the single-unit coefficients of variation of the inter-spike intervals of all cell types being >0.5 (Fig. 2C, left panel) and membrane potential traces displaying a rather marked asynchrony, measured using standard procedures [32].

Because the role of inhibitory neurons is crucial to control the level of firing activity in the columnar model, we analysed the effects of inactivating different interneuron populations. As shown in Fig. S1, the activity of all neurons drastically rose when inhibitory neurons were shut down. In particular, we blocked the transmission of signals from inhibitory to all other neurons, resulting in a sharp increase in pyramidal cell firing rates, which in turn drove the inhibitory firing rate up (even though this firing of inhibitory neurons was not able to suppress the columnar activity).

We controlled for the size of the cortical column model, as it has been reported that not all measurements in spiking columnar models scale linearly with network size and finite-size scaling studies must be performed [30]. When increasing the size of the network in our model and rescaling the weights accordingly (see Methods), we obtained the same firing rate statistics for sufficiently large networks. In Fig. S2, we show the mean firing rates of spontaneous activity using a network of 5k, 10k, 15k up to 80k with similar results in all cases.

Importantly, the dynamics of the model is similar when more biologically realistic aspects are introduced. For example, we may also consider that synaptic weights are lognormally distributed and span several orders of magnitude as reported experimentally [5,33], and the results of the model above do not differ substantially in this case (Supplementary Fig. S3). Likewise, considering a more symmetrical distribution of feedforward input targets (i.e. feedforward input arriving at 25% of excitatory and 25% of PV cells, instead of 25% and 5% respectively) also provide similar results, as long as synaptic weights are properly scaled (Supplementary Fig. S4)

### Three distinct states: spontaneous, feedforward-driven and feedback-driven

We begin by considering three states of the system: spontaneous, feedforward-driven and feedback-driven. In the spontaneous condition (Fig. 1B), the column receives only background input to all cells (Supplementary Table 1), needed to match the cell-type-specific firing rates observed in vivo in mice. When subjected to feedforward input evoking the arrival of bottom-up sensory stimuli (i.e., input to a subset of pyramidal neurons and PV cells in layer 4 (25% of E4 and 5% of PV4), the system displays an increase in neuronal activity in all cell types across all other layers (Fig. 1C). On the other hand, feedback signals corresponding to top-down modulations from higher cortical areas can be simulated by injecting external current into a subset of layer 5 pyramidal neurons [17,19,34,35] and PV cells (25% of E5 and 5% of PV5). Our findings indicate that while feedback input augments the activity of layer 5 pyramidal neurons, it concurrently reduces the firing rates of all other pyramidal neurons in the column, resulting in an overall inhibition of pyramidal cells (Fig. 1D). This phenomenon can be attributed to the modulation of inhibitory neurons, particularly in L4 and L6, which, due to feedback input, increase their activity, inhibiting the pyramidal cells. Consequently, the impact of feedback input stands in contrast to what is observed with feedforward input. To check that no oscillatory regime is present in these conditions (spontaneous, feedforward, feedback) we performed a power spectrum analysis on the mean firing rates and computed the Irregularity of single-unit spike trains quantified by the coefficient of variation of the inter-spike intervals. Our results show (Fig. S5, S6) that no strong oscillatory activity is present in either condition.

### Perturbations of specific cell types in spontaneous and feedforward-driven states

Next, we systematically tested the response of the columnar model to perturbational excitatory inputs to every cell type and layer in the column, for both the spontaneous (Fig. 1B) and stimulus-evoked (Fig. 1C) conditions. Perturbative input targeted to specific cell types allowed us to explore the role of each layer- and type-specific cell in the overall columnar dynamics, as previously done for simpler neural circuits [36]. This will allow us to understand weather the responses to perturbations are dependent on the initial state of the cortical column. To quantify the effects of input perturbations in our network, we defined the response matrix as an array R_XY_ describing the activity change of population X as a result of the perturbative input to population Y. Fig. 3 shows response matrices for spontaneous and feedforward-driven conditions (Fig. 3A and 3B) and their differences (Fig. 3C). We first put the network in a ‘state’, for example spontaneous condition or feedforward condition (i.e. input to a subset of layer 4 pyramidal and PV cells) and then we inject input to one cell group at a time; we then observe the effects on the other neuron groups to build the matrix. Details on how the matrix is computed are provided in the Methods section. We will refer to this process of perturbing one cell group at a time and observing the effect on the others as ‘perturbation analysis’. Overall, effects on any given population are difficult to predict a priori, highlighting the importance of computational tests to guide intuition [36,37]. For example, when we perturbed layer 2/3 pyramidal neurons in the spontaneous state, we observed a substantial increase in layer 6 PV activity (Fig. 3A), even though according to the connectivity data and canonical microcircuit principles [1,23], the strongest anatomical projection from layer 2/3 to layer 5 was between their pyramidal neurons.

**Figure 3.**
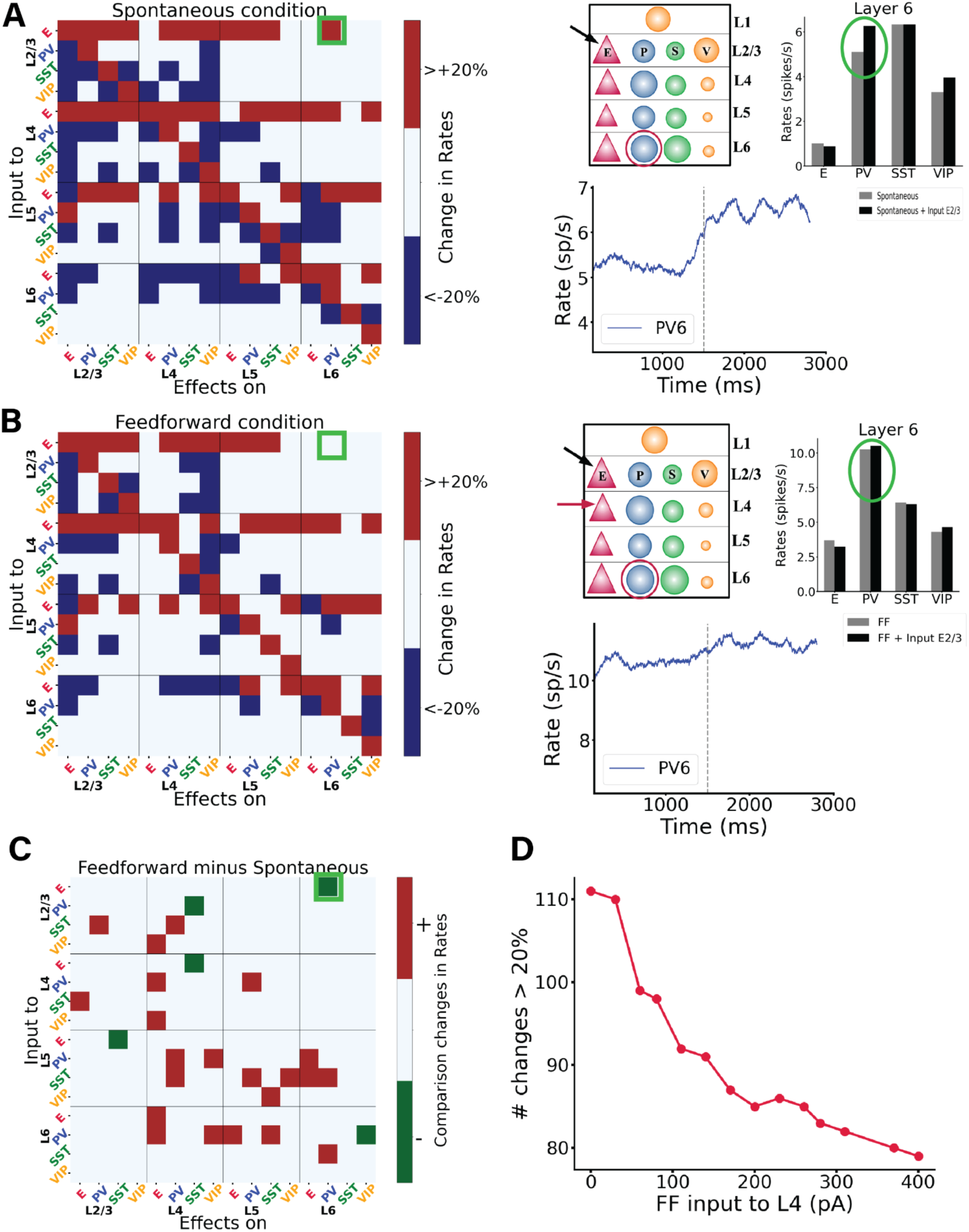
Perturbations of specific cell types in spontaneous and feedforward-driven states. (A) Matrix of input-output relationships of the network in the spontaneous state. We delivered excitatory input to one population (Y-axis; see Methods) and observed its effect on others (X-axis), repeating this for all 16 populations to construct the matrix. Red signifies an increase in mean firing rate by more than 20% above baseline, blue indicates a decrease by more than 20%, and white denotes changes less than 20%. More details on the exact changes in firing rates are shown in Supplementary Fig. S7. We stimulated each subpopulation with a 30 pA DC current and monitored the resultant firing rate changes in all other subpopulations. The right panel of (A) details one specific simulation, where input to the E2/3 population is applied, and the impact on PV cells in layer 6 is measured. The dashed line corresponds to the stimuli onset, the blue trace represents firing rate, while the box plot compares firing rates before and after input injection to E2/3. Here, the PV interneurons in layer 6 respond more robustly than in the feedforward condition depicted below. Note that the firing rate starts to rise a bit before the stimulus onset –this is due to the integration time window of 100 ms used to estimate the firing rates. (B) Displays the response matrix for the feedforward-driven state, wherein excitatory input is provided to a subset of L4 pyramidal cells and PV cells. The right panel presents the same example as in (A) under this condition. (C) Matrix illustrating the difference between the two conditions, where white indicates no change, red a positive difference (i.e., a firing rate increase in the FF condition as compared to spontaneous condition), and green a negative difference (see Methods for detailed specifications). Comparing the two shows that overall, in the feedforward case many firing rate changes become less relevant (<20%) compared to the spontaneous case. (D) shows the number of changes >20% in the corresponding perturbation matrix. We conducted perturbation analysis for 14 different network conditions, defined by varying Feedforward (FF) input to layer 4, and resulting in a 16×16 matrix for each condition, though these matrices are not displayed here. Each condition varied the input strength to a subset of excitatory and PV neurons in layer 4 (25% of E4 and 5% of PV4), values on the X-axis range from 0 to 400 pA. The Y-axis represents the number of alterations >20% (or <-20% in the respective matrix (the sum of red and blue squares). Increased input to layer 4 results in fewer perturbation-induced changes in the firing rates of other populations.

The results also change depending on the condition: as Fig. 3 shows, a small perturbative input to pyramidal neurons in layer 2/3 increases the firing rate of PV cells in layer 6 in spontaneous conditions, but the same stimulus induces only minor changes in PV activity during the stimulus-evoked condition (compare rate plots in 3A and 3B). Other features remain invariant across conditions: perturbing layer 6 pyramidal neurons acts as a suppressor of superficial layers (2/3 and 4) and activator of deep layers (5 and 6) in both conditions. This aligns with experimental observations which characterize layer 6 as a translaminar inhibitor [28]; our model reveals that such inhibition may well be the consequence of complex cortical interactions and not only of the recruitment of local inhibitory cells. The model might help us explain the dynamics causing the increase in activity within layer 5. By observing the connectivity matrix (Fig. 1A), we note that pyramidal cells in layer 6 are strongly connected to PV cells in layer 5, making it intuitive to expect an increase in the activity of PV cells in layer 5 when layer 6 pyramidal cells are perturbed. This, in turn, might be one of the factors responsible for the decrease in activity of pyramidal cells in layer 5. A case-by-case comparison between the effects in spontaneous and feedforward-driven conditions (Fig. 3C) reveals a wide variability of effects across the matrix. For example, our model predicts a difference in the response of SST cells in layer 4 when perturbing pyramidal cells in layer 4 in two scenarios. In the spontaneous case, the SST cells increase their activity by 22%, whereas in the feedforward case, a change of only 6% is induced, in Fig. S7 the exact percentage changes for all perturbative inputs are shown. We can speculate that feedforward input has a stabilizing effect in the context of perturbation. In fact, when comparing the two conditions (Fig. 3C), we note that many perturbations become less marked in the feedforward case (see also Fig. S7).

Other differences between the two scenarios supporting this idea are the following:

1. In the spontaneous case, pyramidal cells in layer 4 show rate changes induced by the perturbation of several interneurons in several layers (VIP 2/3, PV 4, VIP 4, PV 6). In the feedforward case, these changes are smaller (<20%), see Fig. 3C.
2. In the spontaneous case, perturbation of SST cells in layer 5 shows a great (<-20%) negative rate change in many cells in non-superficial layers (PV 4, PV 5, VIP 5, E 6, PV 6); this is not observed in the feedforward condition (Fig. 3C).
3. In the spontaneous case, perturbation of PV cells in layer 6 induces a negative rate change of <-20% in several cell types (E4, VIP 4, E5, SST5), which is less marked in the feedforward condition (Fig. 3C, S7).

This observation shows that in the feedforward case, the model is more robust against small perturbations. Moreover, our model can act as a prediction for experiments. This analysis indicates, as also shown by previous studies [36], that simple intuition sometimes leads to wrong conclusions, or overgeneralization, in models of cortical circuits with multiple cell types.

We next performed an extensive perturbation analysis for 14 different scenarios. In each scenario, we varied the strength of (non-perturbational) feedforward input injected into 25% excitatory neurons and 5% PV cells in layer 4, mimicking activation by retino-thalamic input. The input strength ranged from 0 to 400 pA. For each condition, we generated a perturbation matrix by giving input to one neuronal group at a time and observed the changes in the others (as above). We then counted the number of marked changes (more than a 20% increase or decrease) in their firing rates upon the injection of perturbational input into any population. We observed that networks with stronger feedforward input display less marked changes upon perturbations than networks with weak feedforward input (Fig. 3D). The input condition the number of changes was being in a state of receiving weak (∼30 pA) feedforward input, indicating that input perturbations (presumably coming from other cortical circuits) have a stronger influence on visual cortex when animals receive weak visual input.

This result might seem in contrast with our common intuition: Feedforward (FF) input would generally increase the firing rates of neurons across the network. As a result of increased firing rates, the intuitive expectation is that any perturbations (especially to inhibitory neurons) would likely have a more pronounced effect in reducing the overall firing rates because there’s more activity to suppress. However, contrary to these hypotheses, we observe that the network becomes more resilient to perturbations. We may explain this phenomenon by noting that when feedforward (FF) input is strong, the overall cellular activity reaches such a high level that minor perturbations are insufficient to alter the collective outcome.

### Perturbations in the feedback-driven state

We subsequently examined the model’s response to perturbative excitatory inputs to each cell type and layer in the feedback-driven state, i.e. in a cortical column in which 25% excitatory neurons and 5% PV cells in layer 5 receive feedback drive (Fig. 1C). The corresponding response matrix [36] and its comparison with the spontaneous case are depicted in Fig. 4 (A and B panels respectively), which shows that a majority of perturbations results in positive changes, indicated by the prevalence of red over green squares. This can be explained from the fact that it is easier to excite a network that is already inhibited, in this case by feedback to layer 5. Consequently, in the presence of feedback, perturbations from adjacent circuits (like neighbouring cortical columns) become effectively excitatory overall. This indicates that feedback input to cortical columns may serve as a control mechanism to modulate lateral interactions across V1 columns, as also suggested by experimental findings [38,39]. In Supplementary Fig. S8 we also provide the matrix with the exact percentage changes of firing rates caused by the perturbative inputs.

**Figure 4:**
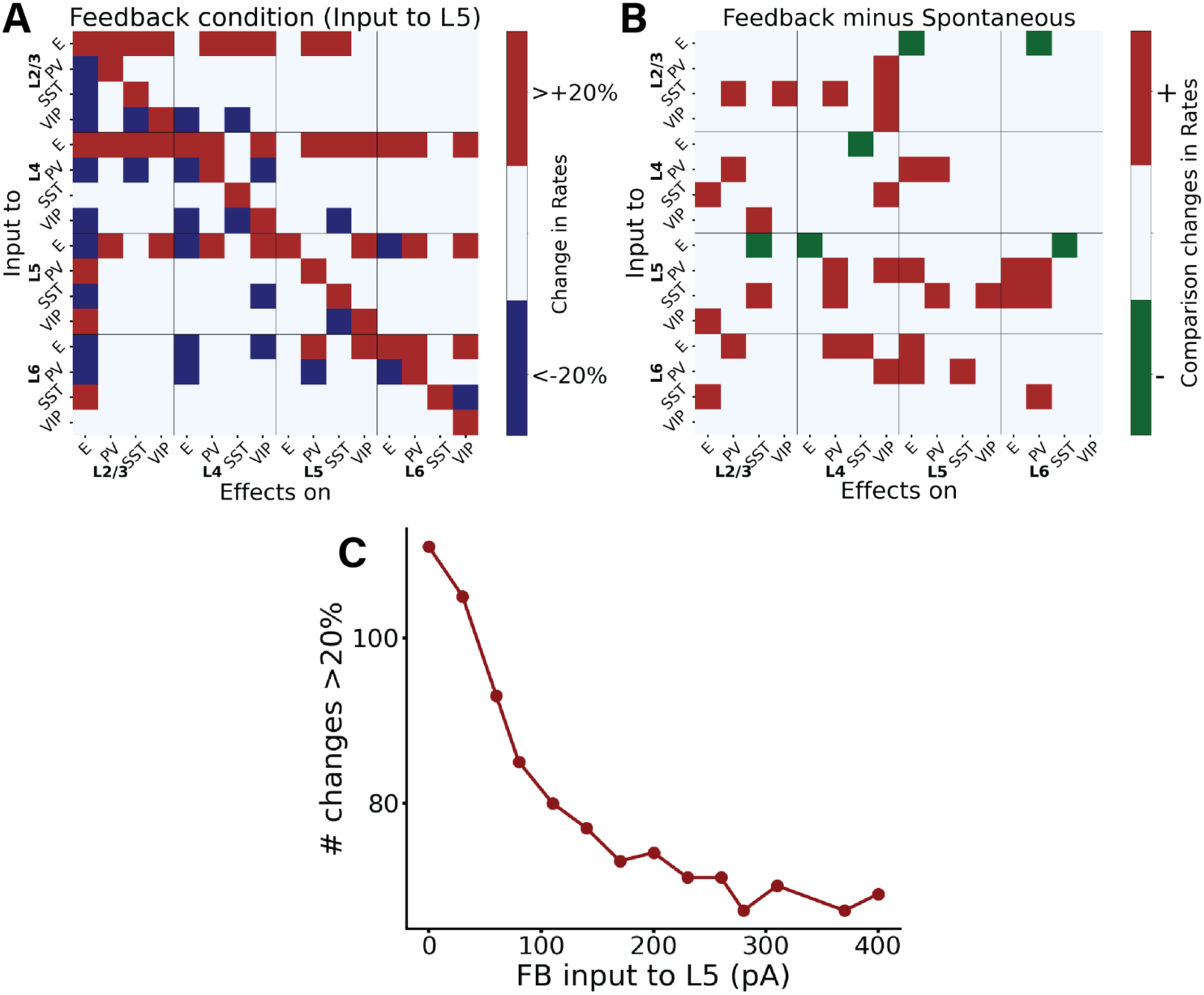
Perturbational input-output matrix in feedback-driven condition. (A) Perturbational matrix of input-output relationships within the network. With feedback input (input to pyramidal cells in layer 5) applied, we administered input to one population (indicated on the Y-axis) and observed the effects on the others (X-axis). This process was repeated for all 16 populations to compile the matrix. Red signifies an increase in the mean firing rate exceeding 20% above the initial value (with only input to L5), blue denotes a decrease greater than 20%, and white represents a change less than 20%. In Supplementary Fig. S8 the exact percentage changes are shown. (B) Comparative matrix between the feedback and spontaneous conditions (layout as in Fig. 3C). (C) shows the number of changes >20% (or <-20%) in the corresponding perturbation matrix. We conducted perturbation analysis for 14 different network conditions, defined by varying Feedback (FB) input to layer 5, and resulting in a 16×16 matrix for each condition, though these matrices are not displayed here. Each condition varied the input strength to a subset of excitatory neurons and PV cells in layer 5 (25% of E5 and 5% of PV5), values on the X-axis range from 0 to 400 pA. The Y-axis represents the number of alterations >20% (or <-20%) in the respective matrix (the sum of red and blue squares). Increased input to layer 5 results in fewer perturbation-induced changes in the firing rates of other populations.

We next performed an extensive perturbation analysis for 14 different scenarios, similarly to what was previously done for the feedforward case. In each scenario, we varied the strength of (non-perturbational) feedback input injected into a subset of excitatory neurons (25%) and PV cells (5%) in layer 5. The input strength ranged from 0 to 400 pA. For each condition, we generated a perturbation matrix by giving input to one neuronal group at a time and observed the changes in the others. We then counted the number of marked changes (more than a 20% increase or decrease) in their firing rates upon the injection of perturbational input into any population. We observed that networks with stronger feedback input display fewer marked changes upon perturbations than networks with weak feedback input (Fig. 4C). In Supplementary Fig. S9, we also provide, among the marked changes, the percentage of positive changes (being roughly half of all of them) for each scenario. This result is similar to the one previously obtained by varying feedforward input strength. Probably the reason behind this is that when we have a strong feedback input which is governing the overall behaviour small perturbative inputs have irrelevant effect.

Contrary to the more stereotypical feedforward input from sensory streams, which commonly targets excitatory neurons and PV cells in layer 4, feedback projections have more diverse cellular targets [40] and synaptic projections may target several layers, with the notable exception of layer 4 which is consistently avoided by feedback projections [17,41]. This makes evaluating the effect of feedback modulation computationally more difficult. Therefore, in addition to the previously analysed feedback condition (input to a subset of pyramidal and PV neurons in layer 5), we explored four alternative feedback configurations (Fig. 5A), in which input respectively arrived at: (i) subset of pyramidal cells and PV cells in layer 2/3 (25% of E2/3 and 5% of PV2/3), (ii)subset of pyramidal cells and PV cells in layers 2/3 and 5 (25% of E2/3, 25

**Figure 5:**
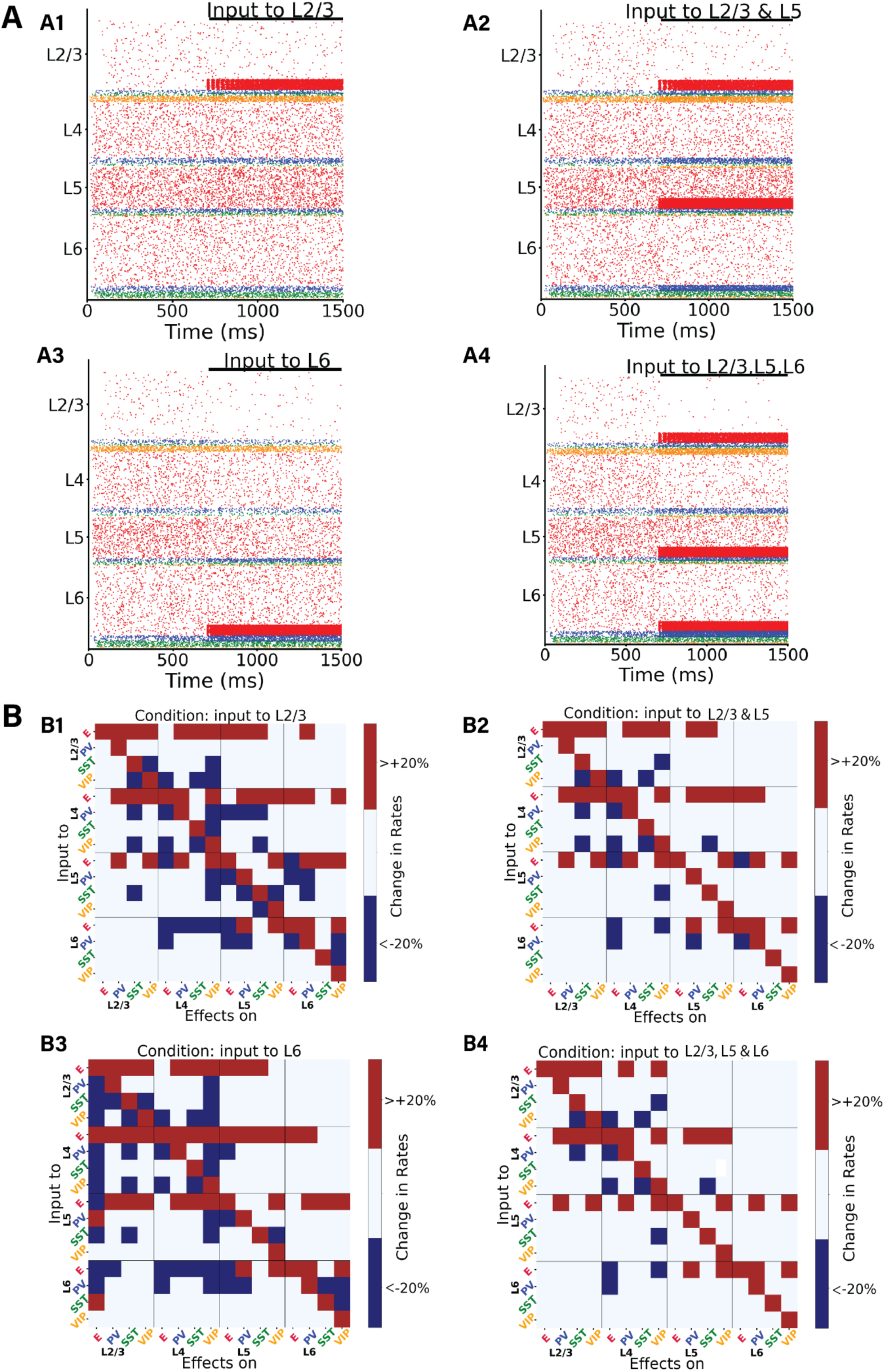
Perturbation analysis in four different feedback-driven states. (A) Raster plots depicting four different states. The network state after 700 ms in the raster plot indicates the condition for the subsequent perturbation analysis. Upon injection of the input in A1-A4) we can observe a synchronisation of the neurons receiving the input, this effect then disappears with time. (B) Perturbation Input-output matrices for four distinct states: B1) Feedback input to subset of E and PV neurons in layer 2/3. B2) Input to a subset of E and PV neurons in layer 2/3 and layer 5. B3) Input to subset of E and PV neurons in layer 6. B4) Input to subset of E and PV neurons in layer 2/3, 5 and 6. Within each condition, we delivered 30 pA DC current to one population (Y-axis) and observed the resultant effects on the others (X-axis). We replicated this procedure for all 16 populations to derive each matrix. For color code, see Fig. 3. Exact changes in percentage firing rates are shown in Supplementary Fig. S11.

% of E5, 5% of PV2/3 and 5% PV5), (iii) subset of pyramidal and PV cells in layer 6 (25% of E6 and 5% of PV6), and (iv) 25% pyramidal cells and 5% PV cells in layers 2/3, 5 and 6. The evoked laminar spiking activity for each of these four alternative configurations (with feedback input arriving at 700 ms) is displayed in Fig. 5 panels A1-4, and we observed overall excitation for case A1 (especially in L2/3 and L5), slight inhibition in case A2 (leaving aside L5 and L2/3 pyramidal cells), overall inhibition in case A3 (leaving aside L6 cells), again in agreement with the translaminar inhibitory effect of layer 6 observed experimentally [28] and overall inhibition in A4 (leaving aside the cells directly receiving the input).

The respective perturbational input-output matrices for configurations A1-4 are shown in Fig. 5B (subpanels B1-B4), and reveal that perturbational effects are overall similar for most feedback configurations, with the number of changes >20% (or <-20%) being approximately equal for all configurations (Suppl. Fig. S10). This suggests that, from a global point of view, responses to perturbational input (such as lateral interactions) will be relatively insensitive to the type of feedback pathway present, as long as the column is in a feedback-driven state. This robustness of feedback modulation effects should, however, be contrasted with several notable exceptions. For example, perturbations to SST cells in layer 6 alone never triggered a response >20% in pyramidal cells of other layers, except for the configuration B3, feedback targeting layer 6. In this scenario, pyramidal cells in layer 2/3 increase their activity when perturbation is applied to layer 6 SST cells, and decrease it when it is applied to layer 6 PV cells –highlighting cell-type selective modulation across layers. Different perturbation effects are observed in layer 2/3 SST populations when layer 5 pyramidal cells are stimulated, varying across configurations. Specifically, SST activities increase >20% in configurations B3, whereas their firing rates changes less drastically B1,2,4 (See also Fig. S11). To quantify the differences between the conditions we used the Frobenius norm (see Methods): we compute the pairwise distances between all the matrices to see which conditions are closer to each other. This result is shown in Supplementary Fig. S11 as well as the matrices with the exact changes in percentage firing rates.

### Integration of feedforward and feedback signals in the column

Under realistic conditions, cortical columns in V1 may operate in a mixed regime, in which the column receives both sensory feedforward and modulatory feedback signals simultaneously. To explore this case, we analysed the impact of feedback modulation (i.e., input to layer 5) on a stimulus-evoked response that had been introduced shortly before (input to layer 4). The two inputs exhibited opposing effects: while feedforward input to layer 4 (introduced at 500 ms) augmented the activity of other pyramidal cells in all other layers, feedback input (introduced at 1100 ms) reduced it (Fig. 6A). In the isolated feedback scenario (Fig. 1C), we established that feedback input led to a decrease in the firing rates of all other pyramidal neurons than those in layer 5, resulting in a comprehensive inhibition of the circuit. Consequently, when both feedforward and feedback inputs were introduced at different intervals, their antagonistic roles became evident. For example, in layer 2/3 (and to some extent in layer 6), the feedback input adjusted the excitatory firing rates to levels similar to those observed prior to the introduction of feedforward input, effectively neutralizing the impact of the latter (Fig. 6A, 6B). To be more specific, upon arrival of feedback input to a subset of layer 5 pyramidal cells and PV cells at 1100 ms, there was a discernible decrease in the activity of pyramidal cells across all layers, naturally with the exception of layer 5. This change can be attributed to inhibitory neurons, especially those in layers 4 and 6. Due to feedback input to layer 5 pyramidal cells projecting to these neurons, they substantially increased their activity across all layers, thereby inhibiting the pyramidal cells in L2/3, L4 and L6.

**Figure 6:**
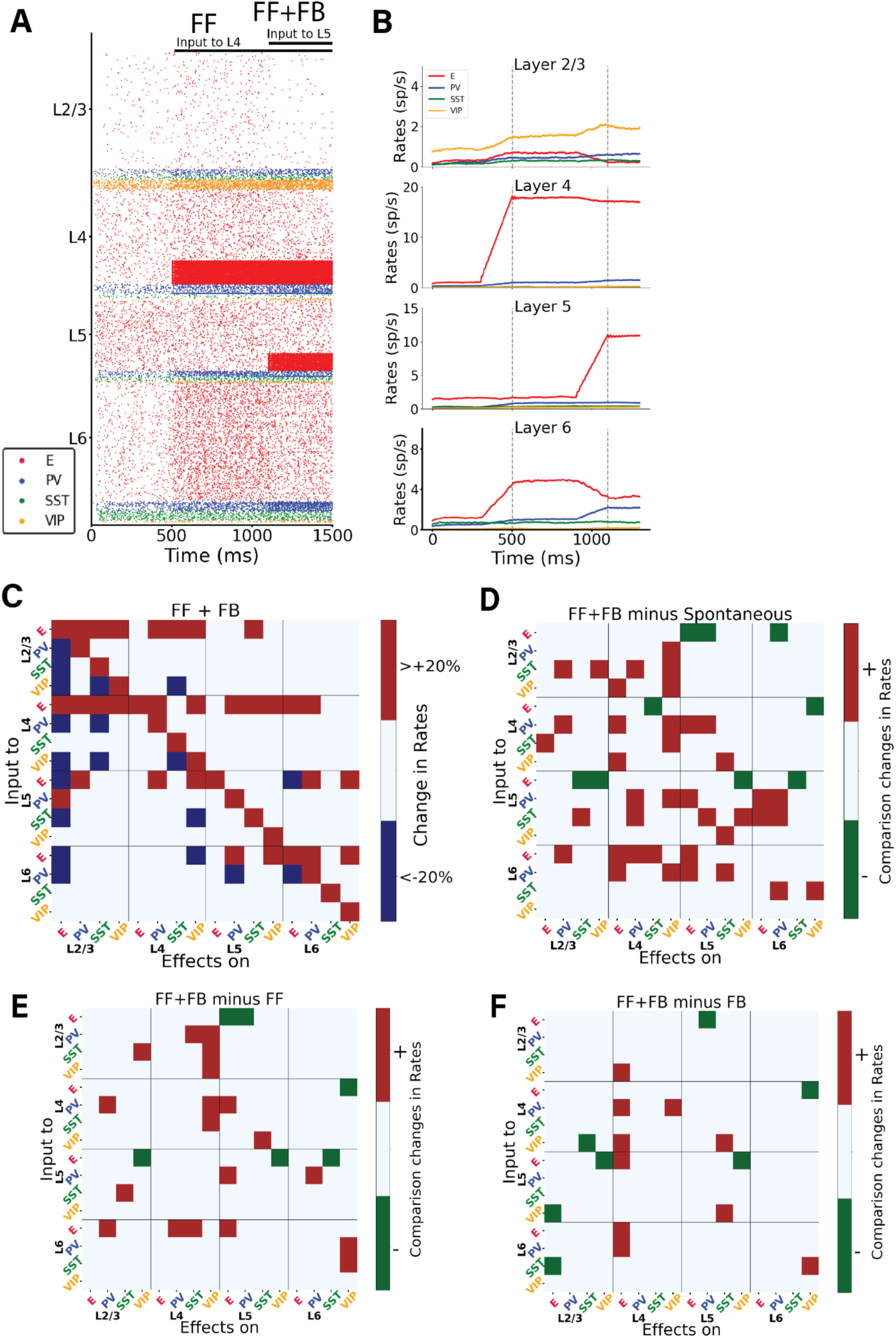
Interaction between bottom-up and top-down signals in the cortical column model. (A) Raster plot of network activity with bottom-up and top-down input. At 500 ms a prolonged (feedforward) input is applied to 25% of layer 4 excitatory neurons and 5% of PV cells. and at 1100 ms an input is applied to 25% layer 5 excitatory neurons and 5% PV cells. The input to L5 has a largely opposite effect compared to the input to L4: it decreases the activity of excitatory neurons in layers 2/3, 4 and 6. (B) Mean firing rate traces of the various populations in each layer. Vertical dashed lines: onset of FF at 500 ms; onset of FB at 1100 ms. Note that firing rates starts to rise a bit before the stimulus onset, due to the integration time window of 100 ms used to estimate the firing rates. (C) Matrix of input-output relationships of the network. When both inputs to L4 and L5 are present, we delivered a perturbative current of 30pA to one population (plotted on the Y-axis) and observed the effect on the others (X-axis). We repeated this procedure for the 16 populations to obtain the matrix. Color code as in Fig 3. (D) Matrix showing the difference between the feedback & feedforward situation and the spontaneous condition. (E) Matrix showing the difference between the feedback & feedforward situation and the feedforward only condition. (F) Matrix showing the difference between the feedback & feedforward situation and the feedback only condition. In Supplementary Fig. S12 the exact percentage changes of firing rates upon injections of the perturbations are shown.

We subsequently conducted a systematic examination of the model’s response to perturbative excitatory inputs directed at each cell type and layer within the column, specifically under conditions where both feedforward and feedback inputs were active (Fig. 6). The response matrix for the columnar model reveals intricate reactions to cell-type specific inputs. In Supplementary Fig. S12 the exact percentage changes are shown. For instance, there is a inter-laminar inhibition affecting all cell types in layers 2/3 and 4 when pyramidal neurons in layer 6 are excited, reflecting again the role of layer 6 as translaminar inhibitor reported experimentally [28,34]. Conversely, inputs to pyramidal neurons in layers 2/3 and 4 induce widespread excitation in almost all cells within those layers. Lastly, stimulating interneurons results in a combination of effects. In Figure 6, panels D, E, and F offer a direct comparison of the response matrices from the spontaneous state (Fig. 3A), the feedforward-driven state (Fig. 3B), and the feedback-driven state (Fig. 4A) to the perturbation matrix derived from the state with combined feedforward and feedback inputs. In the state in which feedforward and feedback drives are combined in the same column, the total number of changes >20% (sum of blue and red squares) is smaller than that in the feedforward-driven or feedback-driven state, (Supplementary Fig. S10).

We then analyzed the case in which feedforward is constantly present, and we put the network in different scenarios varying the feedback input. This resulted in 14 different cases with FB varying from 0 to 400 pA. We conducted a perturbative analysis in each of the states and quantified the number of changes >20% (or <-20%), further categorizing them into percentages of positive and negative alterations. We replicated these experiments with FB input held constant and varied the strength of FF input instead. In both experimental setups, an increase in the strength of the input consistently led to a decrease in the number of changes in firing rates. Detailed results of these analyses are presented in Supplementary Fig. S13.

Lastly, for completeness of our analysis, we conducted new perturbative analysis for some of the network conditions (spontaneous, FF, FB and FF+FB), using different values of perturbative excitatory and inhibitory input to each cell group. We varied the perturbation current in the range (−40, −30, −20, −10, 10, 20, 30, 40) pA. The results are shown in Supplementary Fig. S14. As one would expect, the number of changes increases with the strength of the perturbation (in absolute value) and decreases as we increase the threshold of change with respect to baseline (%). Likewise, the positive and negative branches present slight asymmetries (as the effect of inhibitory perturbations is bounded by the fact that firing rates must remain positive). The irregularity of the neural activity under different stimulation conditions is also shown in Supplementary Figs. S15 and S16 for completeness.

## Discussion

The intricate dynamics of cortical columns, particularly in response to various excitatory inputs, have long been a subject of interest in neuroscience [1,42]. Our study offers a comprehensive exploration of these dynamics in a data-constrained computational model of a V1 cortical column with a realistic connectivity and multiple cell types and postsynaptic receptors, with an emphasis on dynamic cell-to-cell interactions across four different states: spontaneous, feedforward-driven, feedback-driven, and a feedforward-feedback combination [43,44].

Our perturbation analysis serves as a valuable predictive tool for experimental tests. We have shown that the response elicited by stimulating a specific neuronal population can vary significantly based on the network’s initial state, whether it be spontaneous, driven by feedforward (FF) input, feedback (FB) input, or a combination of FF and FB. This variability in response highlights the critical role of the network’s initial conditions in shaping its subsequent dynamics. For example, our model predicts that upon perturbative input to pyramidal cells in layer 2/3, the response of PV cells in layer 6 varies depending on the initial state—being in the spontaneous condition or under feedforward influence. There is a pronounced response in PV cells for the former, while the response becomes negligible for the latter. Furthermore, we demonstrated the stabilizing effects of feedforward inputs: when the network is in an elicited state, it becomes difficult to perturb. As a result, the change in firing rates due to perturbation inputs will be minimal and often <20%, as shown in Fig. 3D. A similar pattern is observed for states with high feedback inputs, as indicated in Fig. 4C.

Another primary finding of our study is the distinctive roles of inhibitory neurons in network dynamics. For instance, our model predicts that in an experimental setting with top-down modulation of V1—such as input to neurons in layer 5—providing stimuli to different inhibitory neurons in layer 6 will induce contrasting responses in the E2/3 neurons, depending on the specific inhibitory subgroup targeted (Fig. 4A and S8). This underscores the specialized functions of various inhibitory cell types in a variety of states. Laminar-specific optogenetic stimulation, which has been developed recently [45], could be used to empirically test these predictions, along with others derived from our model.

Our findings underscore the nuanced, state-dependent interplay between different layers and cell types within the column, in line with previous results on the complex dynamics linked to cell heterogeneity [36,46–49]. The pronounced translaminar inhibition observed when exciting pyramidal neurons in layer 6 (Fig. 5A panel A3), contrasted with the widespread excitation triggered by inputs to pyramidal neurons in layers 2/3 (Fig. 5A panel A1) and layer 4 (Fig. 1B) shows not only the layered specificity of the cortical processing but also the complicated regulatory mechanism in the neuronal network dynamics. The model suggests that excitation of layer 6 pyramidal cells indeed triggers translaminar inhibition, as previously shown experimentally [28]. However, such overall inhibition is not linked to a simple recruitment of local interneurons in each layer as previously suggested, because different interneuron types (in layer 2/3 and 4) also undergo a decrease in activity. Instead, our model suggests that layer 6-triggered translaminar inhibition is a network effect that involves non-trivial interactions across different layers and cell types.

The mixed effects observed upon stimulating (SST, VIP and PV) interneurons suggests the importance of considering the diverse roles of different neuronal populations within the column [1,5,10,50,51]. For instance, our perturbation analysis of feedback states (Fig. 4,5, S8, S11) indicates contrasting effects when perturbing different interneurons types. In contrast to PV cells, perturbations to SST cells in layer 6 alone never triggered a response >20% in pyramidal cells of other layers, with the exception of the configuration B3, feedback targeting layer 6). In this particular set up, layer 2/3 pyramidal cells increase their activity >20% when perturbation is applied to layer 6 SST cells, and decrease it when PV cells were targeted– highlighting cell-type selective modulation across layers. Such a pattern aligns with the classical disinhibitory pathway [48] observed here in a trans-laminarly fashion: elevated SST activity in layer 6 leads to decreased activity in layer 2/3 pyramidal cells. However, this disinhibitory effect was less marked (<20%) in most other states (see Fig. S11), except in the scenario with feedback to layer 5 (Fig. 4A). This reiterates the critical role of initial network states in interpreting the outcomes of perturbations.

Our model suggests that the dynamics between pyramidal, SST and VIP cells are more complex that what is usually assumed when envisaging a classical disinhibitory pathway [48]. Indeed, SST-mediated disinhibition of pyramidal cells was less marked in our model for most of the states or configurations considered, suggesting that the occurrence of disinhibition in real circuits is likely driven by context-dependent plasticity mechanisms, which aligns with previous experimental and computational results [36,52,53].

The effect of feedforward and feedback inputs, both individually and in combination, generate different responses. While feedforward inputs generally augmented activity of pyramidal cells across all layers (Fig. 1C), feedback inputs often had an inhibitory effect (Fig. 1D), which became especially evident in layer 6 (Fig. 5-A3). This observation aligns with the notion that feedback mechanisms often serve to modulate lateral inter-columnar interactions (as observed experimentally [38,39]) or fine-tune cortical responses [54], potentially to optimize information processing or to prevent runaway excitation [55]. Interestingly, the competing effects of feedforward and feedback inputs, when introduced simultaneously, revealed the delicate balance maintained within the columnar circuitry [19,31]. The ability of feedback inputs to L5 to effectively neutralize the impact of feedforward inputs (Fig. 6A, 6B), especially in layer 6, suggests a robust regulatory mechanism at play: inputs to L5 increase the activity of inhibitory neurons in other layers, such as layer 6, which in turn reduces the activity of pyramidal cells, thereby mitigating the initial increase caused by feedforward inputs. Such dynamics might be crucial for ensuring that the column operates within an optimal regime, avoiding both hypo- and hyperactivity.

### Limitations

#### Stimulus-evoked responses

While our cortical column model provides a reasonable match to existing experimental data for spontaneous conditions, its performance in other states such as FF- or FB-driven must be properly compared to experimental observations in future studies. In particular, stimulus-evoked firing rates of around 15 spikes/s in layers 4 (for FF-driven conditions) and 5 (for FF-driven conditions) are higher than typical rates of 5∼7 spikes/s observed in electrophysiological recordings [23,56] and closer to values of ∼10 spikes/s provided by other models [29]. This does not affect the conclusions of our study, as we have shown that our main findings remain valid for weaker FF and FB inputs (Figs. 3D, 4C and S13). Considering strong FF and FB input also provides a better idea of how our model responds to a broader input range, which might be useful when adapting this model to other brain regions with higher stimulus-evoked responses. Finally, evoked firing rates of 15 or even 30 spikes/s are more reasonable when considering responses to preferred orientation rates [56], and models aiming for more accurate predictions regarding stimulus-evoked responses should consider incorporating stimulus tuning and selectivity properties.

#### Synaptic plasticity

A further factor which plays an important role in generating rich neuronal dynamics is synaptic plasticity, which we have not incorporated in the model for the purposes of this work. Processes such as short-term synaptic plasticity and spike-time dependent plasticity have been shown to result in complex behaviors such as neuronal oscillations [28]. Synaptic plasticity and neuronal oscillations are critical for learning, memory, and coordinated neural activity. Further iterations of the model could incorporate these elements to investigate how long-term changes in synaptic efficacy and oscillatory patterns might influence the function and stability of the cortical column within the various conditions analyzed in this work. Moreover, exploring perturbations in such plastic models would provide a deeper understanding of neural dynamics. Nonetheless, the perturbations responses in our current model, even without incorporating synaptic plasticity, hints at how such plasticity might modulate working memory and learning processes. If the column shows increased sensitivity to perturbations under certain conditions, it may suggest a heightened capacity for synaptic changes at the basis of learning and memory formation.

#### Multi-compartmental neurons

In this work, we employed a leaky integrate-and-fire (LIF) neuron model. Introducing multi-compartmental neurons would add significant biological detail, offering a richer exploration of biophysical network dynamics. A single-compartment model, as used here, restricts our capacity to simulate the intricate intracellular processes occurring within actual neuronal structures. Multi-compartment models could simulate dendritic processing and back-propagating action potentials, offering a more detailed understanding of how individual neurons contribute to overall columnar activity. Such an expansion could enhance our comprehension of local processing within neurons and how activity propagates through the network.

#### Thalamus and higher order structures

Our model’s structure also lacks the thalamus; thalamic input is approximated as currents targeting layer 4, omitting direct modelling of the thalamus and the V1-LGN feedback loops that are integral to visual processing and attention mechanisms. Nevertheless, the simulation of external inputs to layer 4 neurons in our simulations may mimic the effects of thalamocortical loops. Incorporating thalamic interactions would enhance the model by simulating more accurate sensory input dynamics and enabling the exploration of attentional modulation on cortical column function. Future models could include thalamic inputs to study their effects on cortical computations, which is particularly relevant to conditions like schizophrenia, where thalamocortical interactions are frequently disrupted [57].

The absence of higher-order structures is another limitation; feedback from higher areas is modelled as currents targeting different layers but a comprehensive study of the interactions with higher visual areas is not included. Additionally, the model does not currently simulate natural stimuli, which could drastically change the responses of the cortical column due to the complexity and variability inherent in real-world inputs. However, such an investigation was beyond the scope of this study.

### Implications of the model

A recent study emphasize the unique role of layer 5 (L5) neurons in subcortical loops, which participate in feedback mechanism into the cortex and thalamus [1,54]. Our analysis highlights the importance of L5 in counterbalancing feedforward excitation and shows the significance of this layer in both FF and FB processing. Given the special interest in L5 cells, our model can serve as a predictive tool for the effects of perturbations on this layer and help in designing relevant experiment, thereby affirming the functional significance of this layer. We can speculate that the modulation of activity in L5 cells may have far-reaching implications for the tuning of cortical-thalamic interactions, potentially affecting sensory processing and cognitive functions associated with these pathways.

The perturbation approach utilized in our study could be adopted more broadly as an analytical tool in other computational models, such as those exploring predictive coding in the visual cortex [58]. For example, this approach could benefit models that incorporate interneurons, to dissect the individual and collective contributions of interneurons to network behavior, thus enhancing our understanding of their roles in cortical computations.

Our model reveals cascading effects across various neuronal types, which presents a challenge for decoding specific roles of these cells in predictive coding [59] (PC) frameworks. Identifying which neurons are responsible for coding errors or predictions becomes complex when considering the multitude of interactions in the complex dynamics of our cortical column model. However, this complexity is reflective of the nature of cortical computations and suggests that understanding coding in such networks may require an approach which looks at emergent properties rather than reductionist approaches. The current model can suggest experiments that may be conducted within simplified, rate-based models. Such models could be better suited for this exploration and to determine which cell types encode errors or predictions.

Lastly, our findings reveal a stabilizing effect of feedforward (FF) input, hinting that cortical columns may be inherently constructed to provide stability and resilience against perturbations. This resilience could be a natural consequence of the need for reliable information processing in the face of constantly changing environmental inputs. Our observations could inform hypotheses regarding the evolution of neural architectures in the cortex, optimized for maintaining stability while allowing for the flexibility required for adaptive behavior.

#### Relevance to Cortical Tasks

Our findings may have implications for tasks involving the cortex, such as working memory and contextual learning. The E/I balance, central to our cortical column model, is crucial for the proper functioning of cortical circuits involved in these cognitive functions. Disruptions in this balance have been linked to cognitive deficits and neuropsychiatric disorders. All interneuron types in the model contribute to maintaining a balanced network, and the state-dependent effects offer insights into how the cortex might adapt to cognitive tasks. For instance, tasks involving working memory often require information manipulation based on the current cognitive context, and our model suggests that cortical responses to such tasks may vary significantly with the underlying state of neural activity. Integrating feedforward and feedback information is crucial for complex cognitive tasks, and our findings on the distinct effects of these input types may relate to how the cortex integrates sensory information with higher-order cognitive processes during contextual learning.

## Conclusion

Previous research has highlighted the challenges of predicting activity changes in cortical circuits based on simple intuitions [60]. The multifaceted interactions between different cell types, layers, and input modalities necessitate a detailed computational approach to grasp the underlying mechanisms. In conclusion, our findings provide valuable insights into the operational intricacies of cortical columns, emphasizing the importance of both feedforward and feedback-induced state changes in shaping neural responses.

The role of PV cells and NMDA receptors in modulating E/I balance points to their potential involvement in the pathology of neurological disorders where this balance is disrupted such as schizophrenia and autism. Our results highlight the delicate equilibrium required for proper cognitive functioning, and suggest that disruptions in this balance may contribute to the cognitive and behavioral disturbances characteristic of these conditions. Future studies might delve deeper into the specific molecular and biophysical mechanisms that underlie these dynamics, potentially paving the way for targeted therapeutic interventions in disorders marked by such neural imbalances.

## Methods

### Model architecture

The cortical column model, shown in Fig. 1, is composed of a total number (N_total_) of 5,000 neurons. The model consists of four cortical layers each containing pyramidal neurons, PV, SST and VIP cells (layers 2/3, 4, 5, 6) and one layer containing only VIP cells (layer 1). Each of the four ‘complete’ layers harboured pyramidal neurons (85% of cells) and inhibitory interneurons (15%; each inhibitory group in a given layer was represented by a particular percentage out of this 15%). The fractional sizes of neuron types in each layer (of the total number in the column) and those of each cell type in each layer were taken from the Allen institute database [23] and are reported in Supplementary Tables 2 and 3. All neurons received background noise modelled as coming from the rest of the brain (Fig. 1). The levels of background noise that each type of cell received can be found in Supplementary Table 1, the tuned spontaneous firing rate effectuated by the Poissonian spike generators underlying noise generation connected to each group differed amongst cell types.

We use the term *connection* with reference to subpopulations or groups, defined by the pre- and postsynaptic neuron types and the layer they are in. The connection probability defines the probability for each possible pair of pre- and postsynaptic neurons to form a connection between them. If *p*=0.1 this connects all neuron pairs of the two specific groups with a probability of 10%. The connectivity probability matrix *P* is defined by the 16×16 + 16×2 = 288 connection probabilities *p* between the 17 considered cell groups (4 types in each of the 4 layers plus 1 group in layer 1; a group thus potentially receives inputs from the other 16 cell groups and also potentially projects to all of them). The connection probability matrix *P* used to constrain the model is available in the portal of the Allen Institute database (https://portal.brain-map.org/explore/models/mv1-all-layers). Each connection has also a particular strength which differs per neuron group. Thus, the strength was specified at the level of neuron type X projecting to neuron type Y. The strengths of connections between neurons are constrained using the matrix *S* available at https://portal.brainmap.org/explore/models/mv1-all-layers. How we set the strengths of single synapse using the matrix *S* is explained below (Eq. 11).

We illustrate how any two groups (X and Y) are connected with an example: VIP cells in layer 1 (X) that have a connection going to SST cells in layer 4 (Y) will all have the same connection strength. However, not all VIP cells in layer 1 are connected to SST cells in layer 4 because they connect with probability *p*.

### Neuron Model

All pyramidal cells and all three types of interneurons are modelled as leaky integrate-and-fire neurons. Each type of cell is characterized by its own, individual set of parameters: a resting potential *V_rest_*, a firing threshold *V_th_*, a membrane capacitance *C_m_*, a membrane leak conductance g_L_ and a refractory period *τ_ref_*. The corresponding membrane time constant is *τ_m_= C_m_*/g_L_. The membrane potential *V(t)* of a cell is given by:

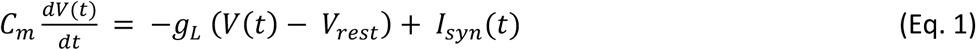

where *I_syn_(t)* represents the total synaptic current flowing into the cell.

At each time point of the simulation, a neuron integrates the total incoming current *I_syn_(t)* to update its membrane potential *V(t)*. When the threshold *V_th_* is reached a spike is generated, followed by an instantaneous reset of the membrane potential to the resting membrane potential *V_res_*_t_. Then, for a refractory period *τ_ref_*, the membrane potential stays at its resting value *V_rest_* and no spikes can be generated. After *τ_ref_* has passed, the membrane potential can be updated again (see Supplementary Tables 5-9 for corresponding parameter values).

### Model of synapses

Each cell group in each layer is connected to all the other groups of the cortical column with its own synaptic strength and probability. The values in matrix *P* indicate the probability that a neuron in group A (e.g. the PV cells in layer 4) is connected to a neuron in group B (e.g. the SST cells in layer 5). Excitatory postsynaptic currents (EPSCs) have two components mediated by AMPA and NMDA receptors, respectively. Inhibitory postsynaptic currents (IPSCs) are mediated by GABA_A_ receptors.

The inputs to model neurons consist of three main components: background noise, external (e.g. sensory) input and recurrent input from within the column. EPSCs/IPSCs due to background noise are mediated in the model exclusively by AMPA receptors (*I_bg,AMPA_(t)*) and those due to external stimuli (i.e. originating from outside the column) are represented by *I_ext_(t)*. This external current may include, for example, constant feedforward or feedback inputs from other brain areas, or brief input perturbations to specific cell types. The recurrent input from within the column is given by the sum of I*_AMPA_(t), I_NMDA_(t), I_GABA_(t).* These are all the inputs from all the other (presynaptic) neurons projecting to the neuron under consideration.

The total synaptic current that each neuron receives is given by:

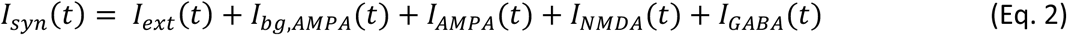

with the last four terms given by

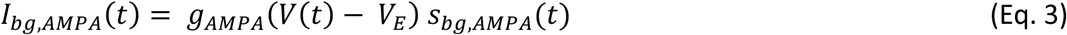

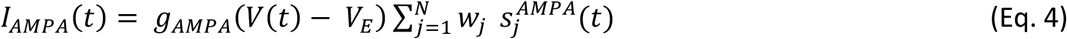

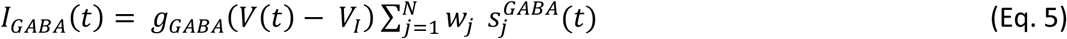

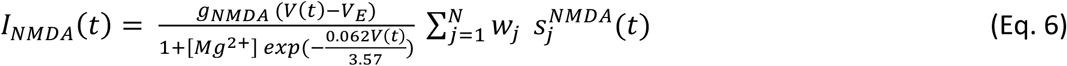

where the reversal potentials are *V_E_*=0 mV, *V_I_ = V_rest_*, and each group of inhibitory interneurons has its own *V_rest_*. The *g* terms represent the conversion factors for the conductance of specific receptor types, and are set to 1 nS. The weights *w_j_* represent the (dimensionless) strength of each synapse received by the neuron. The sum runs over all presynaptic neurons *j* projecting to the neuron under consideration. NMDAR currents have a voltage dependence controlled by extracellular magnesium concentration [61], *[Mg^2+^]=*1 mM. The *s* terms represent the gating variables, or fraction of open channels and their behaviour is governed by the following equations. The amplitudes of EPSCs and IPSCs in our model depend on multiple factors (including the previous spiking history and the value of the membrane potential) and therefore they are not associated exclusively with w_j_ or any other parameter in isolation. Detailed measures of EPSCs and IPSCs for each cell type and layer should be able to help improve the model in the future.

First, the AMPAR channels are described by

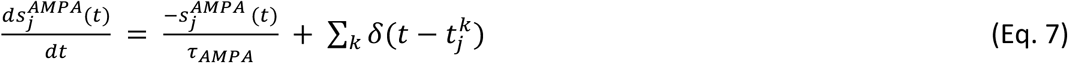

where the decay time constant of the AMPA currents is τ_AMPA_= 2 ms, and the sum over *k* represents the contribution of all spikes (each of which is indicated by delta, *δ*) emitted by presynaptic neuron *j.* In the case of background AMPA currents (Eq. 3), the spikes are emitted according to a Poisson process with rate υ_bg_. Each group of cells in each layer is receiving a different Poisson rate of background noise (see Supplementary Table 1). υ_bg_ is cell-specific, hence each has a specific firing pattern.

The gating of single NMDAR channels is described by

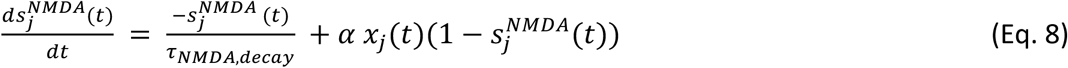

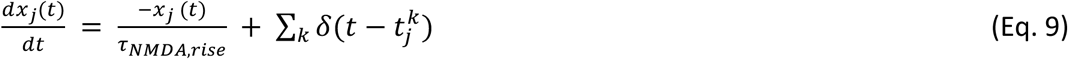

where the decay and rise time constants of NMDAR current are 80 ms and 2 ms respectively, and the constant α=0.5 ms^−1^. The GABA_A_ receptor synaptic variable is described by

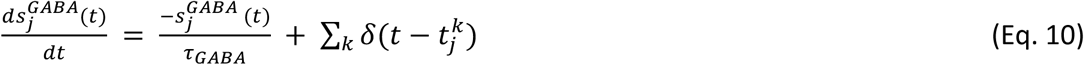

where the decay time constant of GABA_A_ receptor current is 5 ms.

### Parameters of the model

As previously mentioned, each type of cell in each layer is characterized by its own set of parameters: a resting potential *V_rest_,* a firing threshold *V_th_*, a membrane capacitance *C_m_*, a membrane leak conductance g*_L_* and a refractory period *τ_ref_*. The parameter setting s were taken from the Allen Institute database (https://portal.brain-map.org/explore/models/mv1-all-layers). In particular, for each type of cell in each layer the Allen database proposes different subsets of cells (e.g. two different PV subsets in L4), each with his own set of parameters. To simplify the model, we only used one set of parameters for each cell type in each layer, choosing the set of parameters of the most prevalent subset of cells. The parameters *C_m,_ g_L,_ τ_ref_, V_rest_, V_th_* used for each cell type in each layer are reported in Supplementary Tables 5-9.

### Details on background noise

All neurons received background noise, representing the influence of the ‘rest of the brain’ on the modelled area, as shown in Figure 1. Excitatory postsynaptic currents (EPSCs) due to background noise are exclusively mediated in the model by AMPA receptors, denoted as *I_bg,AMPA_(t)* (Eq. 3). The levels of background noise that each group of cells received can be found in Supplementary Table 1. The firing rate υ_bg_ of the background Poissonian pulse generators connected to each group differed among each cell type. Each neuron in every group was connected to its own background Poisson generator. Thus, even though the rates υ_bg_ of the Poisson generators were the same for all neurons within the same group, each specific cell received its own specific pulse train.

### Details on synaptic weights

The weight of each synapse w_j_ from neurons of group A to neurons in group B was set as follows:

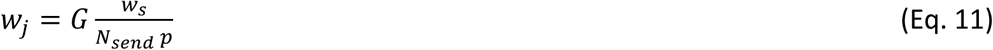

where *G* is the global coupling factor, *w*_*s*_ is the overall strength (measured in mV) between the two connected groups of cells, *N_send_* is the number of neurons in the sending population A, and *p* the probability of connection between the neurons of the two groups (A and B) taken from the experimental probability matrix *P*. For each pair of connected groups, *w*_*s*_ is taken from the experimental synaptic connectivity matrix *S*, defined by the 16×16 + 16×2 = 288 synaptic strengths between the 17 considered cell types (4 groups in each of the 4 layers + 1 group in layer 1). Matrices *S* and *P* can be found at https://portal.brain-map.org/explore/models/mv1-all-layers.> The normalization in Eq. 11 above guarantees that the dynamics and equilibrium points of the system scale properly with the size of the network (Fig. S2). Unless specified otherwise, we choose G=5 mV^−1^, which provides an overall strength of interactions sufficient to make the input from recurrent connections relevant for the spiking activity of individual neurons, but not strong enough to lead to pathological states of over-synchronization.

The spikes generated by a presynaptic excitatory neuron can target AMPA or/and NMDA receptors of the postsynaptic neuron. The AMPA and NMDA receptors are chosen to be present at the excitatory synapse in a 0.8 and 0.2 ratio respectively. Thus, the probability of connection *p* between the neurons of the two groups (A and B) is multiplied by 0.2 for NMDA receptors and 0.8 for AMPA receptors. For example, suppose that excitatory neurons in group A are connected to neurons in group B with a probability *p* (taken from the matrix P). Then the excitatory connections targeting the AMPA receptors of group B will be chosen with *p_AMPA_* = *p*\*0.8 and those targeting NMDA receptors will be chosen with *p_NMDA_* = *p*\*0.2.

All differential equations were numerically solved with a time step of 0.1 ms.

### Probability correction

Since, for realistic cell densities, the connection probabilities in Billeh et al. correspond to a column with radius of r=75 um and our own network of 5,000 neurons would correspond to a column of radius ∼125um, we translated those quantities to better approximate values of uniform connection probabilities.

We performed the translation for each connection type using the correct value of the Gaussian width of each connection, denoted as sigma. These values of sigma are reported in supplementary material of Billeh et al. and also in Supplementary Table 10.

The new correct uniform probability used now in the model are given by

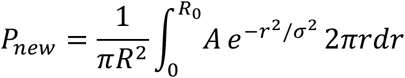

where 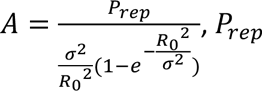 is the probability used previously, R = 125 um, *R*_0_ = 75 um, and sigmas are reported in Supplementary Table 10 along with the result of the calculation.

### Perturbation matrix

We analysed the effects on neuronal populations following the administration of a specific input. Each square in a matrix (e.g. figure 3A) corresponds to an individual simulation. The y-axis specifies the population receiving the input, while the x-axis indicates the observed effect on a different population. To create the matrices, we injected a 30 pA constant current into each group and recorded the ensuing effects on all other neuronal groups. The percentage change in average firing rate, used to construct these matrices, was determined by comparing the mean firing rates after stimulation with the respective baseline rates (prior to input injection). To evaluate the baseline rates we used a time window of 3 seconds of simulation, the same time window was used to evaluate the rates after the injected current. We thus calculated the mean firing rate for each population both before and after the stimuli were introduced. In the matrices, red signifies an increase in firing rate by 20% or more relative to the baseline, blue indicates a decrease in firing rate by 20% or more, and white reflects a change in firing rate that is less than 20% (whether a decrease or an increase). We also provided all the matrices with the exact changes of percentage firing rates in Supplementary figures S7, S8, S11, S12.

It is important to understand that these ‘baselines’ for firing rates (initial states) varied depending on the scenario under analysis: Spontaneous, FF input (with different strength), FB input, or a combination of the two. We always compared the firing rates after the perturbation stimuli with the firing rates prior the input injection, which was specific for each initial state analysed.

To put the model in the desired condition (FF, FB, FF+FB etc) we inject the external input to the neurons through the I_ext_(t) variable in Eq. 2. For example, for the FF condition we inject 150 pA to 25% of excitatory neurons in layer 4 (E4) and 5% of PV cells in layer 4 (PV4). For the FB condition we inject 150 pA to 25% of E5 and 5% of PV5. All the external inputs for the different feedback conditions analysed are targeting a subset of E cells (25%) and PV cells (5%) in the indicated layer and 150 pA constant current is used.

Although feedforward and feedback input are ultimately conveyed to the network via synaptic inputs, the use of constant injection currents is justified by the diffusion approximation. Briefly, spikes arriving at a given neuron from many presynaptic AMPA synapses can be approximated using a constant I_ext_ current in the equation of its membrane potential. Each input contributes a small amount to the overall membrane potential, and the combined effect of these numerous, small synaptic inputs can be approximated by a continuous process (like a diffusion process) when the number of inputs is large. Mathematically, this is akin to a form of the central limit theorem, where the sum of many small, independent random variables (the synaptic inputs) approaches a Gaussian distribution.

### Random seed

When performing the simulations, we made sure to use the same random seed for each run. This method allows us to control the impact of variability, guarantee reproducibility of each trial, and ensure that the changes in the cells’ firing rates from one simulation to the other were due to the perturbation under examination and not to other fluctuations or variables.

### Comparison Matrix

The comparison matrix displayed in Fig. 3C was constructed as follows: a white square indicates no change in the firing rate of a specific population when comparing the spontaneous case to e.g. the FF case. This can occur when the two perturbation matrices we are comparing had both a red squares, both a blue squares or both a white squares (Fig. 3A-3B).

A red square represents one of the following scenarios:

- A transition from a non-marked alteration (<20% or <-20% in absolute terms) to a marked increase (>20%).
- A transition from a marked decrease (>-20% in absolute terms) to a non-marked alteration (<20% or <-20% in absolute terms).
- A transition from a marked decrease (< −20%) to a marked increase (>20%).

A green square indicates one of the following conditions:

- A transition from a marked increase (>20%) to a marked decrease (> −20% in absolute terms).
- A transition from a marked increase (>20%) to a non-marked alteration (<20%).
- A transition from a non-marked alteration (>-20%) to a marked decrease (> −20%).

We also provide the matrices with the exact comparison of percentage firing rates changes in Supplementary figures S7 (panel C), S8 (panel B), S12 (panel B, C, D). Those matrices illustrate the difference in percentage change between the two matrices A and B (i.e. if in matrix A the percentage change is +50% and in B is +30%, the “comparison matrix” will show −20%). The black contour square indicates when a change in sign between the two matrices occurred.

### Matrix Similarity and Frobenius Norm

In this study, the Frobenius norm was used to quantify the similarity between matrices representing the perturbation outcomes of the different conditions (e.g., spontaneous activity, input to L4, L5, etc). The Frobenius norm is a measure used to quantify the difference between two matrices. For a given matrix A, the Frobenius norm is defined as the square root of the sum of the absolute squares of its elements:

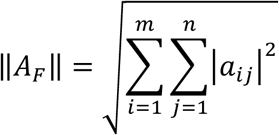

where *a_i,j_* represents the element in the i-th row and j-th column of matrix A, and m and n are the dimensions of the matrix. In the context of comparing two matrices A and B, the Frobenius norm of their difference ||*A_F_*|| provides a scalar value that represents the “distance” between them in terms of their elements. A smaller Frobenius norm indicates greater similarity between the matrices, while a larger norm suggests they are more different.

We computed the Frobenius norm for all pairs of matrices to determine how similar or different they are from one another. The results were stored in a symmetric matrix where each entry (i,j) represents the Frobenius norm between matrix i and matrix j. The heatmap is shown in Fig. S11. In the heatmap, darker colors represent lower norms (indicating greater similarity), and lighter colours represent higher norms (indicating greater difference).

## Acknowledgments

This work was done with the support of EBRAINS and HBP computing services. We thank Matthias Brucklacher, Kwangjun Lee and Parva Alavian for constructive discussions, and Hans Ekkehard Plesser, Sacha van Albada, Walter Senn and Sander Bohte for their useful feedback on early iterations of the work.

## Author contributions

JFM, GM, CP conceived and designed the study; GM and RAD performed the research; GM, RAD and JFM analyzed the results; GM, CP and JFM wrote the manuscript.

## Data and materials availability

All information needed to reproduce the results of this manuscript is available in the main text, Methods section and Supporting information. The Python code developed will be made available upon publication of this work.

## Supplementary figures

**Figure S1:**
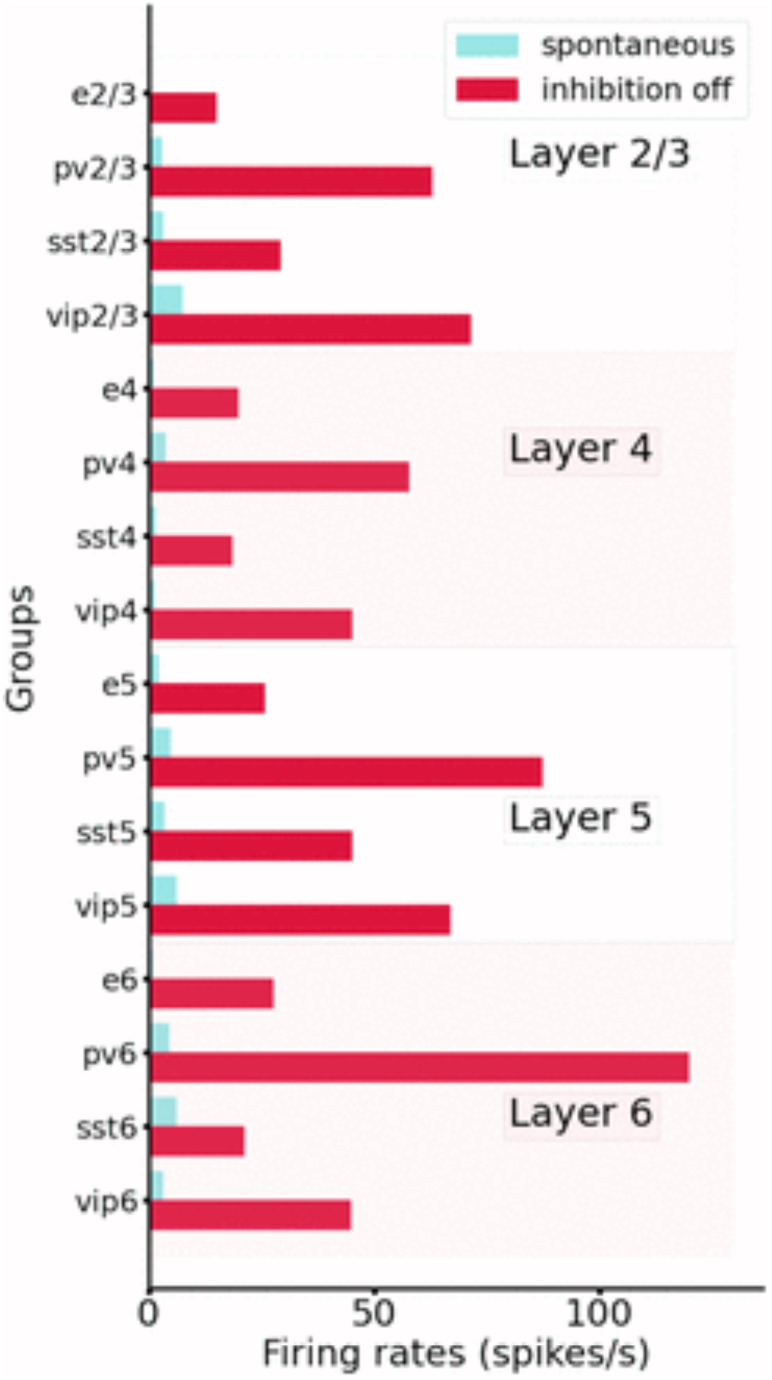
Comparison between spontaneous activity in normal conditions (control, blue) vs. the condition of disabling the output from all inhibitory neurons (while maintaining their capacity to spike). All groups substantially elevated their firing rates, particularly those in deep layers.

**Figure S2:**
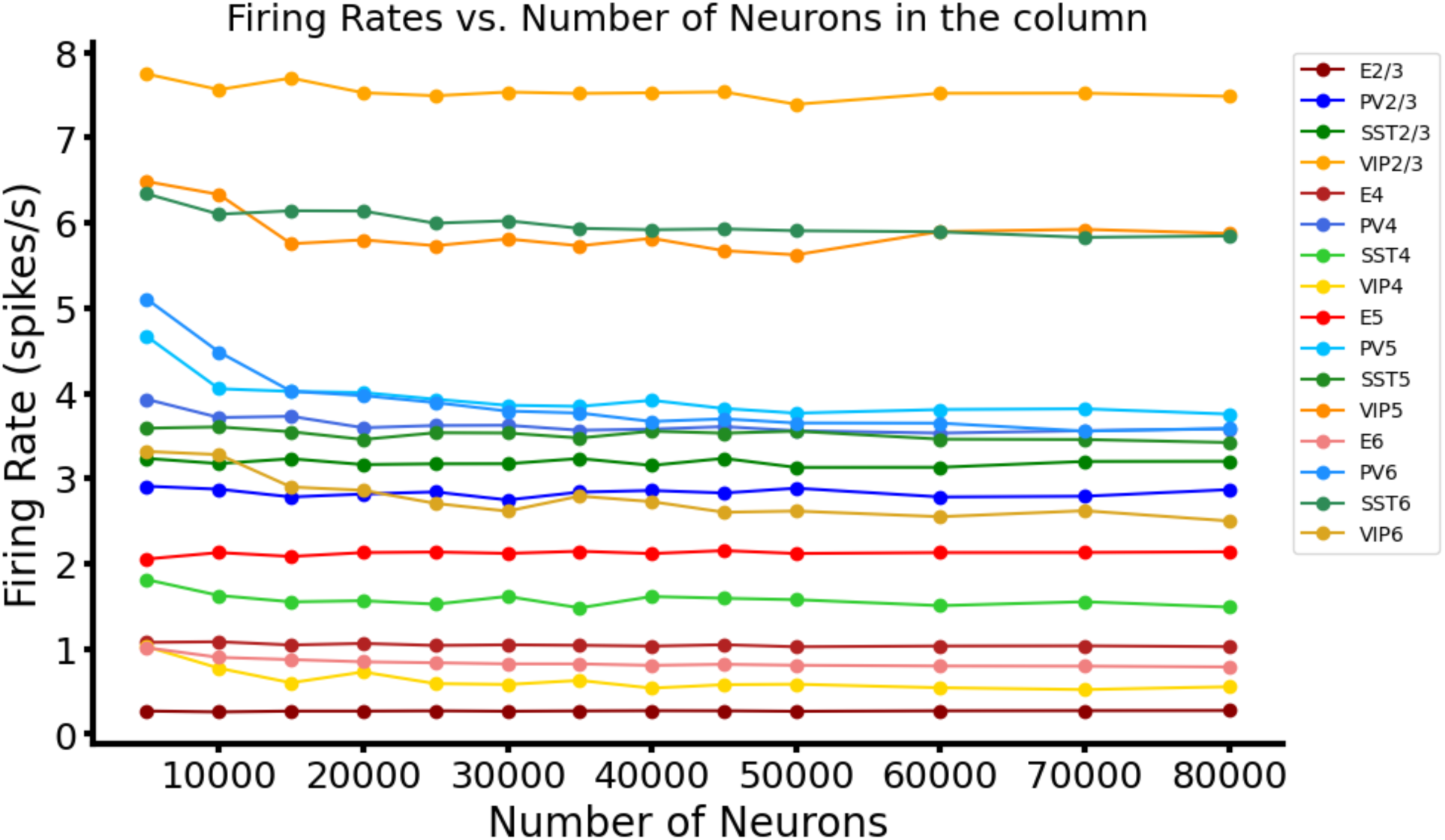
Mean firing rates of all groups for networks of different sizes. We show that, with proper scaling of the weights (see Methods), results for 5k,10k up to 80k neurons lead to very similar results.

**Figure S3:**
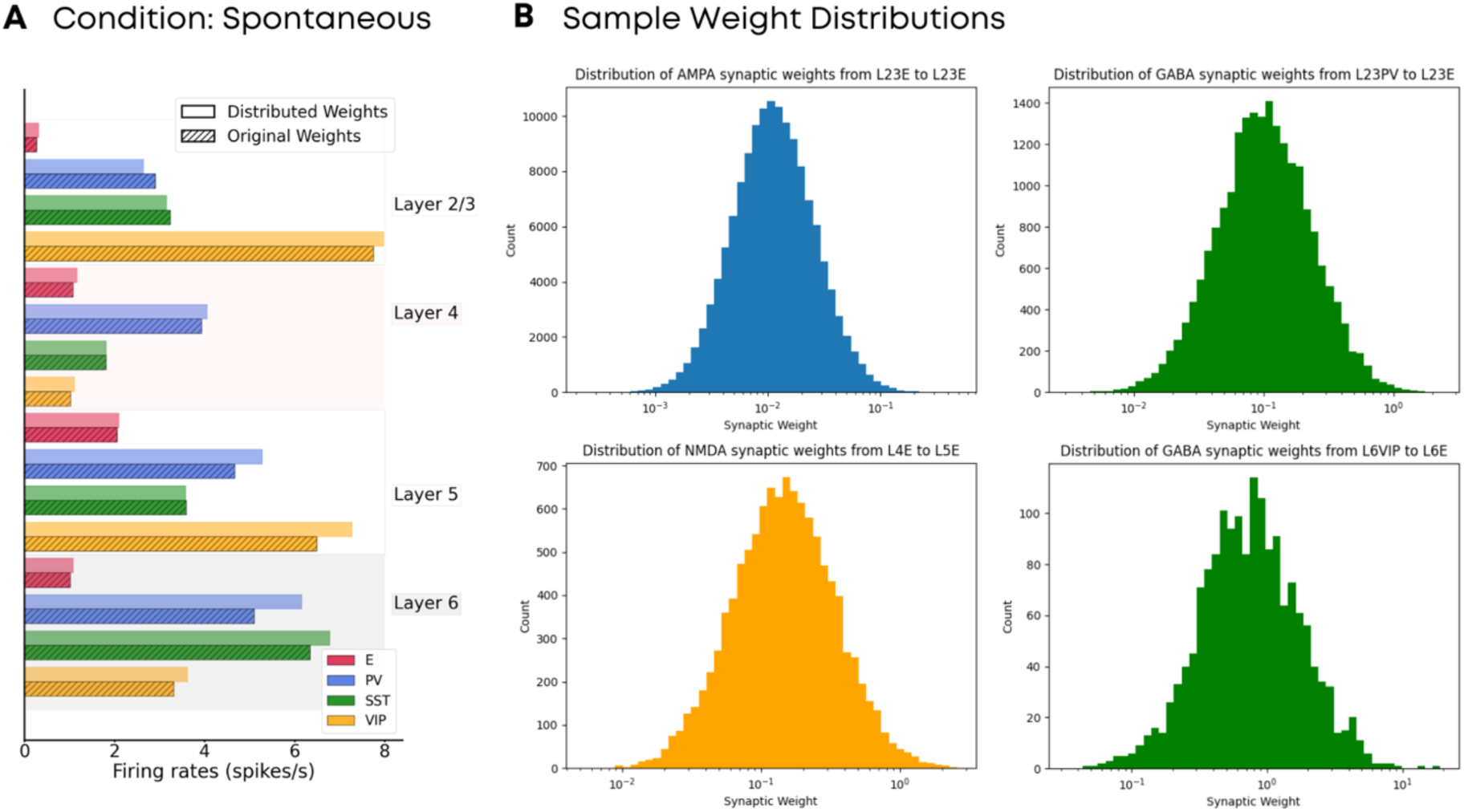
Mean firing rates in the cortical column model, for the spontaneous state, with broad distribution in synaptic weights. A) Comparison between the mean firing rates for all layers and cell types of the original model shown in Figs. 1 and 2 and a model in which values for each individual synaptic weight are chosen from a lognormal distribution, as reported experimentally. The mean of the distribution corresponds to the original value of the projection in the original model, and the standard deviation is set equal to the mean. B) Examples for the resulting distribution of synaptic weights in the modified model, for several synapse and cell types as indicated in the panel titles. For most cases, the distribution of weights spans about two orders of magnitude, in agreement with experimental observations.

**Figure S4:**
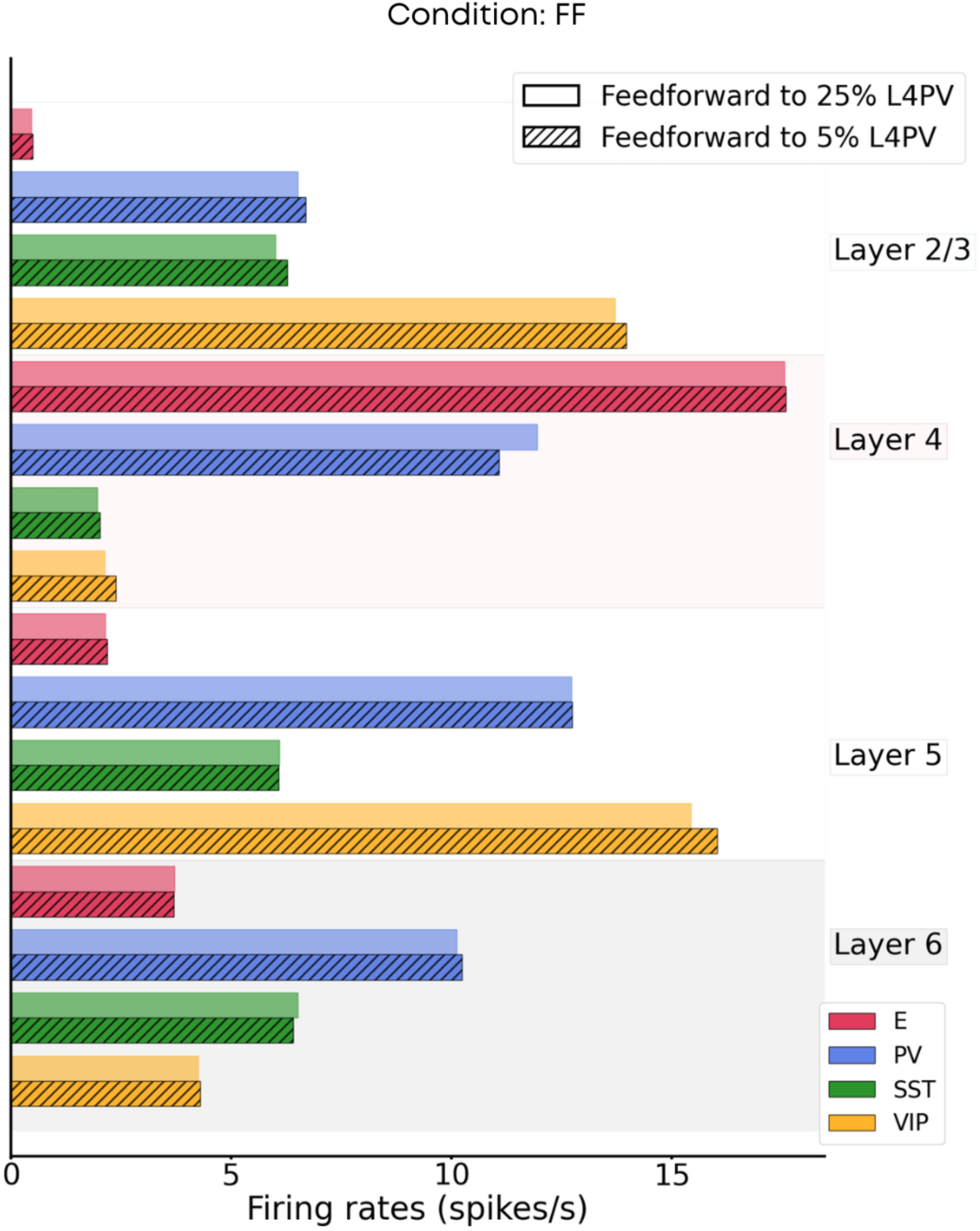
Activity of the cortical column for different targets of the feedforward input. The figure compares the mean firing rates across layers and cell types for the original model shown in Figs. 1 and 2, and a modified model in which the feedforward input arrives at 25% of the PV cells in layer 4, instead of just 5% as in the original model. For the modified model, input weights to layer 4 PV cells were scaled down by a factor of 0.25 to compensate for the additional inhibition.

**Figure S5:**
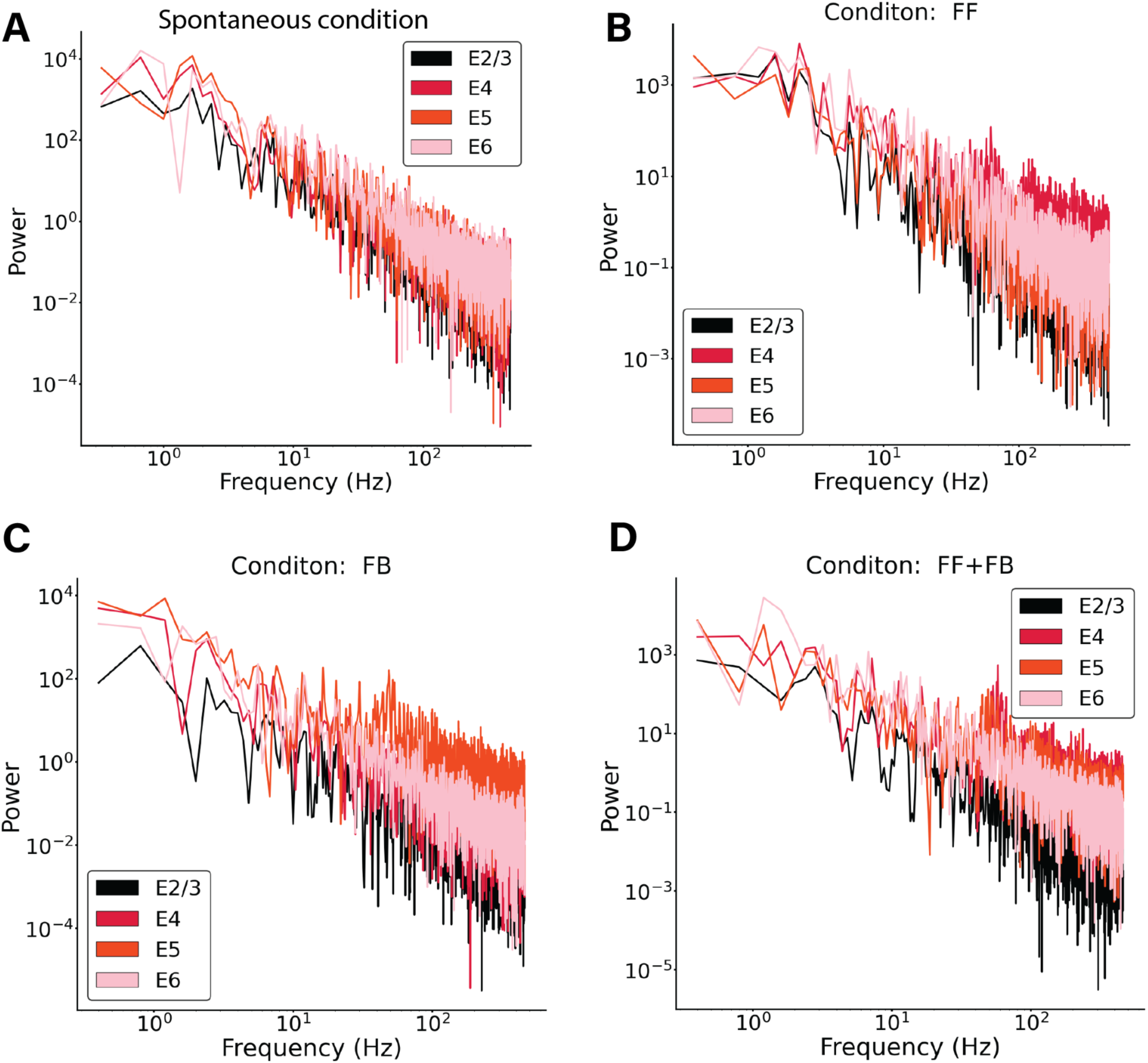
Power spectrum of excitatory mean firing rates across all layers in 4 different network condition (A)-(D). A) Spontaneous condition. B) Feedforward input of 150pA injected in 25% excitatory cells and 5% Pv cells in layer 4. C) Feedback input of 150pA injected in 25% excitatory cells and 5% Pv cells in layer 5. D) Feedback input of 150pA injected in 25% excitatory cells and 5% Pv cells in layer 5 and layer 4. In all conditions no signs of oscillatory activity are shown. The subset of cells receiving the input shows some level of synchronisation, which is reflected in a small peak in their corresponding power spectrum.

**Figure S6:**
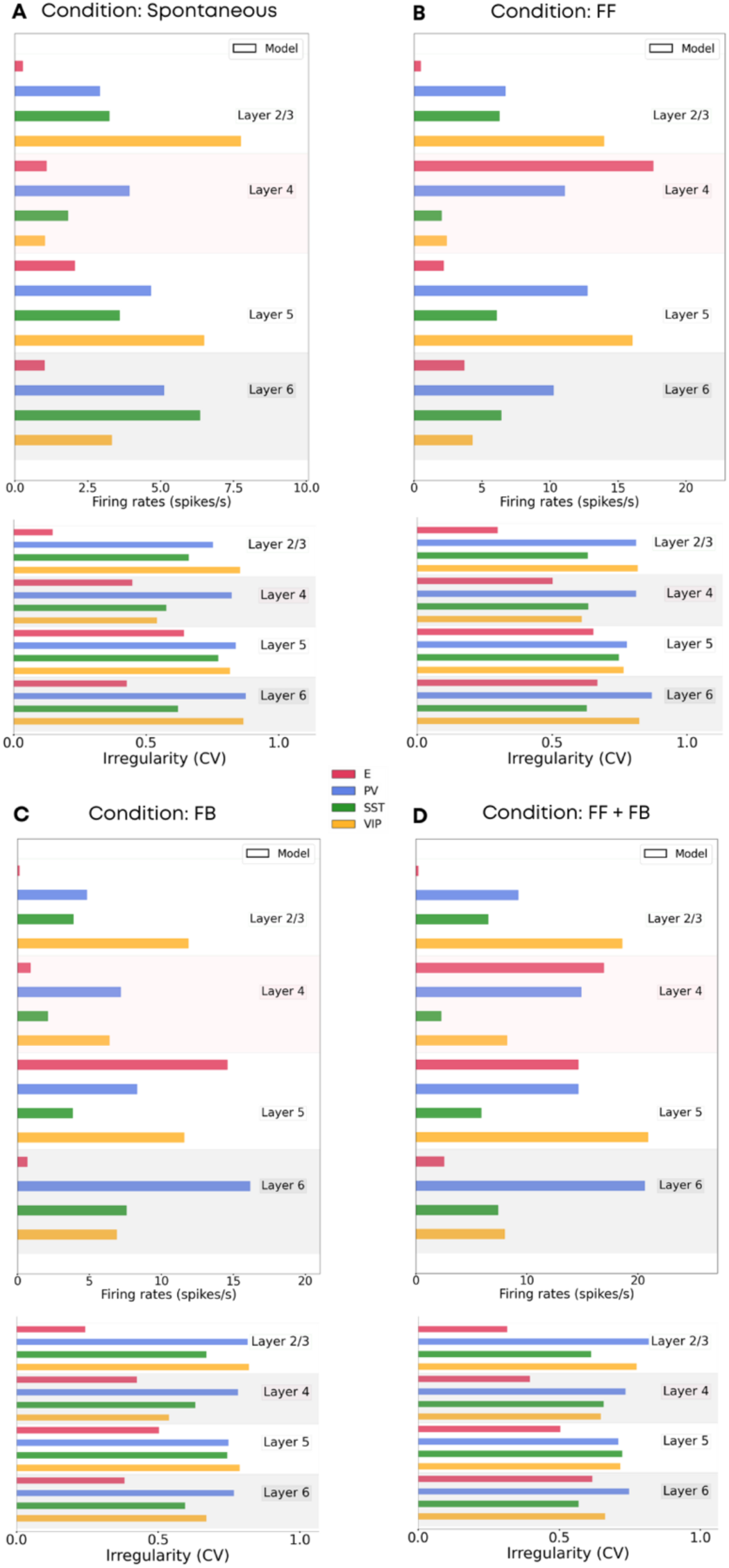
Mean firing rates and irregularity of single-unit spike trains quantified by the coefficient of variation of the inter-spike intervals in four different network conditions. A) Spontaneous condition B) Feedforward input of 150pA injected in 25% excitatory cells and 5% Pv cells in layer 4. C) Feedback input of 150pA injected in 25% excitatory cells and 5% Pv cells in layer 5. D) Feedback input of 150pA injected in 25% excitatory cells and 5% Pv cells in layer 5 and layer 4. In all conditions most cells have a CV >0.5 showing no synchrony. In A) and C) excitatory cells in layer 2/3 have a low CV, probably due to the fact that the firing rate activity is very low, therefore the CV is harder to properly evaluate.

**Figure S7:**
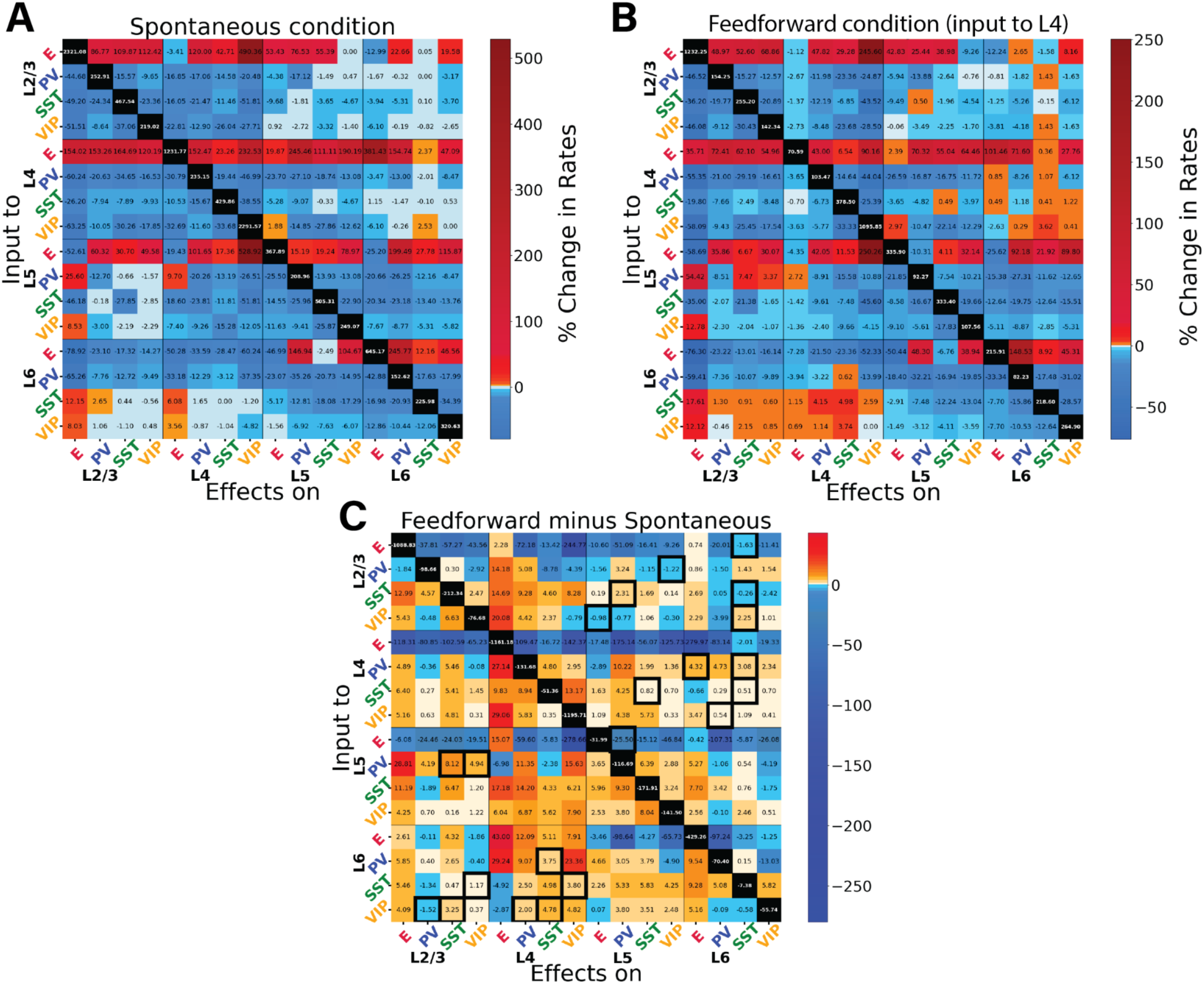
Perturbations of specific cell types in spontaneous and feedforward-driven states. Panels (A), (B) and (C) provide additional information on the data presented in Fig. 3. (A) Matrix of input-output relationships of the network in the spontaneous state. We delivered excitatory input to one population (Y-axis) and observed its effect on others (X-axis), repeating this for all 16 populations to construct the matrix. We stimulated each subpopulation with a 30 pA DC current and monitored the resultant firing rate changes in all other subpopulations. This matrix shows the exact percentage changes of the firing rates compared to the situation pre-perturbative injection. The diagonal (change of firing rate of the perturbed cells) are removed from the colour code and set to black. (B) Displays the response matrix for the feedforward-driven state, wherein excitatory input is provided to a subset of L4 pyramidal cells and PV cells (Input of 150pA to 25% of E4 and 5% of PV4). (C) Matrix illustrating the difference in percentage change between the two conditions (i.e. if in the spontaneous case the percentage change is +50% and in the FF is +30%, the difference matrix will show −20%). The black contour square indicates when a change in sign between the two matrices occurred.

**Figure S8:**
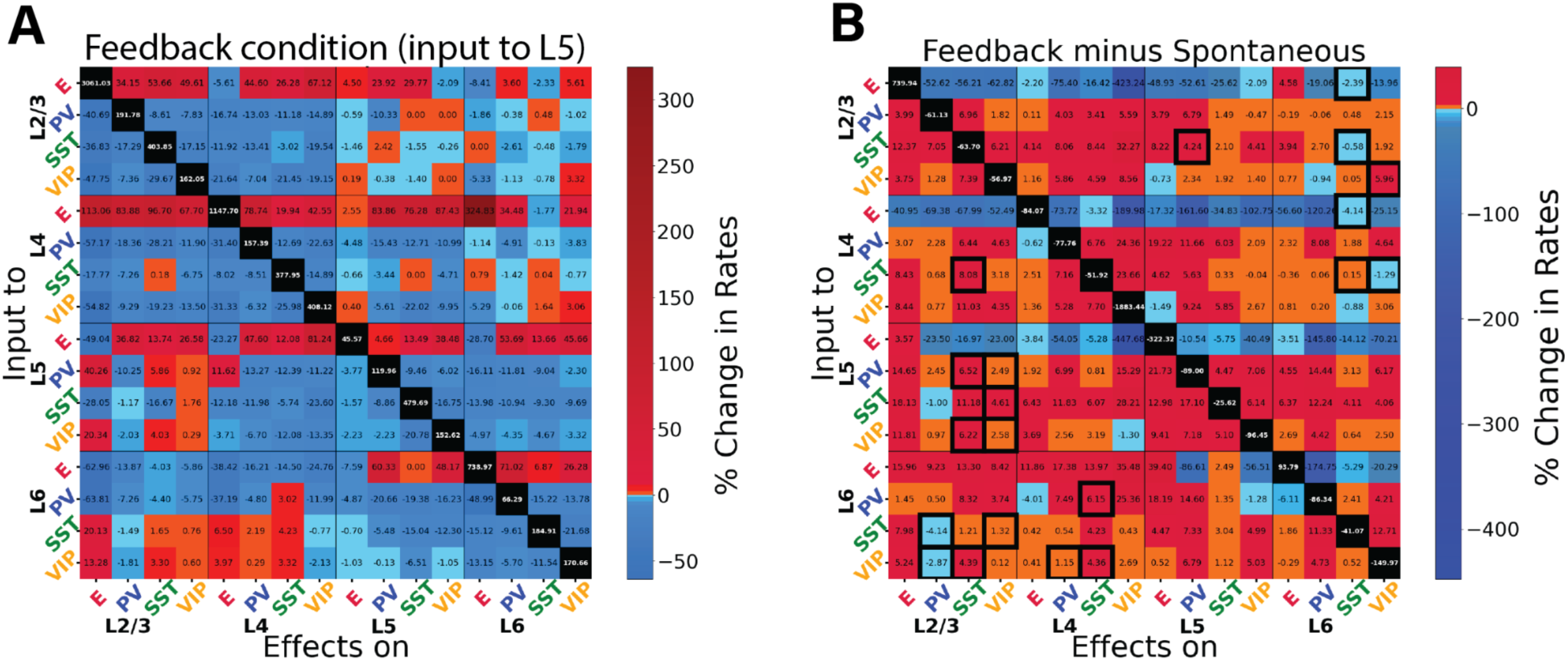
Perturbational input-output matrix in feedback-driven condition. Panels (A) and (B) provide additional information on the data presented in Fig. 4. (A) Perturbational matrix of input-output relationships within the network. With feedback input (150pA input to 25% pyramidal cells and 5% PV cells in layer 5) applied, we administered input to one population (indicated on the Y-axis) and observed the effects on the others (X-axis). This process was repeated for all 16 populations to compile the matrix. We stimulated each subpopulation with a 30 pA DC current and monitored the resultant firing rate changes in all other populations. The matrix shows the exact percentage changes of the firing rates compared to the situation pre-perturbation injection. (B) Comparative matrix between the feedback and spontaneous conditions.

**Figure S9:**
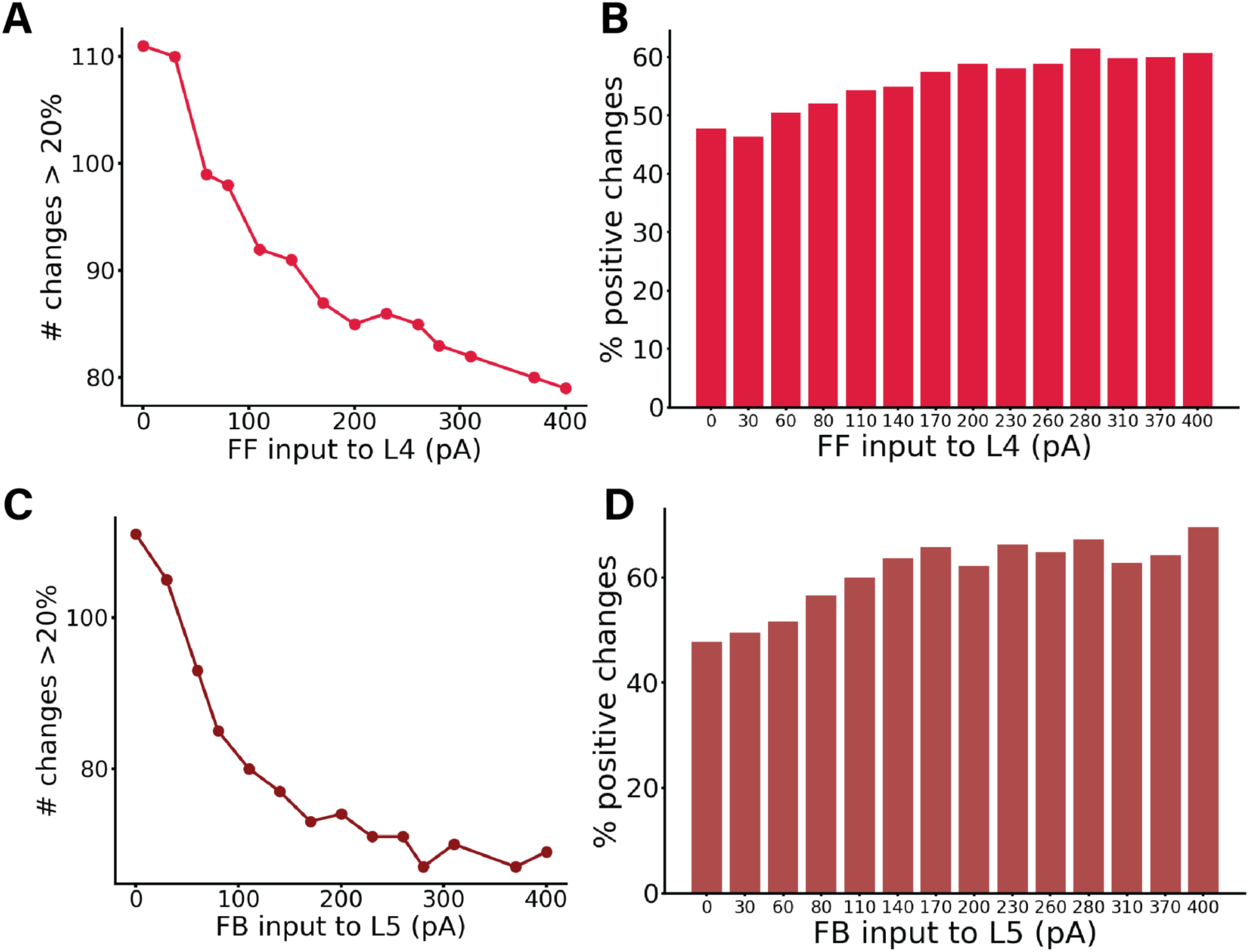
Panels (B) and (D) provide additional information on the data presented in Fig. 3D and Fig. 3C, respectively (shown again here as (A) and (C)). (A) displays the number of changes >20% (or <-20%) in the corresponding perturbation matrix. We conducted perturbation analyses for 14 different network conditions, defined by varying feedforward (FF) input to layer 4, resulting in a 16×16 matrix for each cell group, although these matrices are not displayed here. Each condition varied the input strength to excitatory neurons in layer 4, with values on the X-axis ranging from 0 to 400 pA. The Y-axis represents the number of alterations >20% (or <-20%) in the respective matrix (the sum of red and blue squares). Increased input to layer 4 results in fewer perturbation-induced changes in the firing rates of other populations. (B) shows percentages of positive changes in firing rates elicited by all possible perturbations in each network state, computed from the total number of changes presented in A. The complementary percentages in the panel also provide data for the negative changes. (C) displays the number of changes >20% (or <-20%) in the corresponding perturbation matrix for 14 different network conditions, defined by varying feedback (FB) input to layer 5, resulting in a 16×16 matrix for each cell group, although these matrices are not displayed here. Each condition varied the input strength to excitatory neurons in layer 5, with values on the X-axis ranging from 0 to 400 pA. The Y-axis represents the number of substantial alterations >20% (or <-20%) in the respective matrix (the sum of red and blue squares). Increased input to layer 5 results in fewer perturbation-induced changes in the firing rates of other populations. (D) presents percentages of positive changes in firing rates elicited by all possible perturbations in each network state, computed from the total number of changes shown in C. The complementary percentages in the panel also provide data for the negative changes.

**Figure S10:**
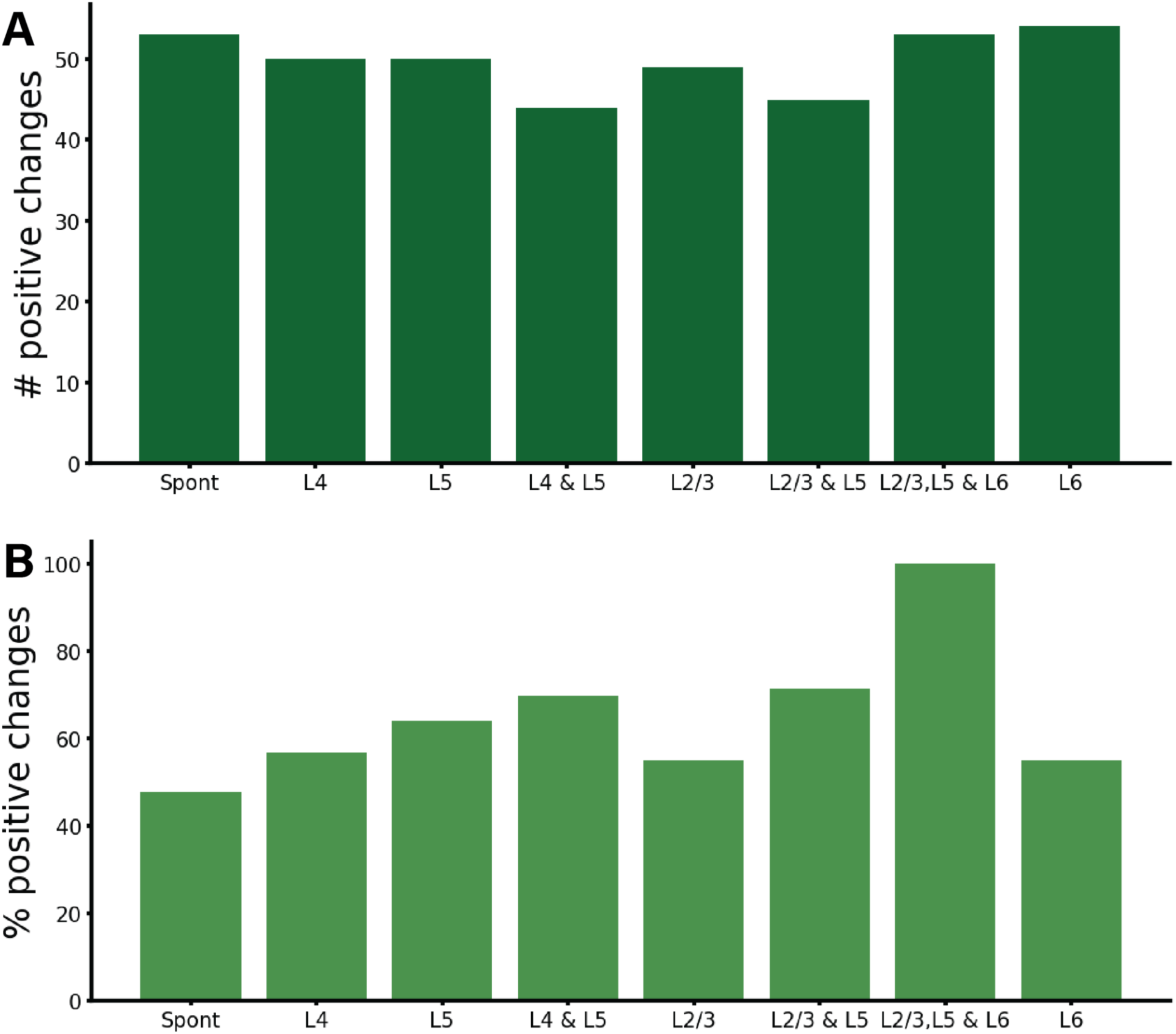
Number of changes >20% (or <-20%) (A) and percentages (B) for positive changes in firing rates elicited by all possible perturbations in the spontaneous state (Spont), feedforward-driven state (L4), feedforward-feedback combination (L4 & L5), and feedback-driven state in all its five possible configurations (i.e. targeting subset of pyramidal and PV in layer 5 (L5),subset of pyramidal and PV cells in layer 2/3 (L23), subset of pyramidal and PB cells in layer 2/3 and 5 (L2/3 & L5), subset of pyramidal and PV cells in layer 2/3,5 and 6 (L2/3, L5 & L6), and subset of pyramidal and PV cells in layer 6(L6)). The complementary numbers for the bottom panel also provide the data for the negative changes. This figure provides extra analysis on the data presented in Fig. 3A, 3B, 4A, 5B, 6C.

**Figure S11:**
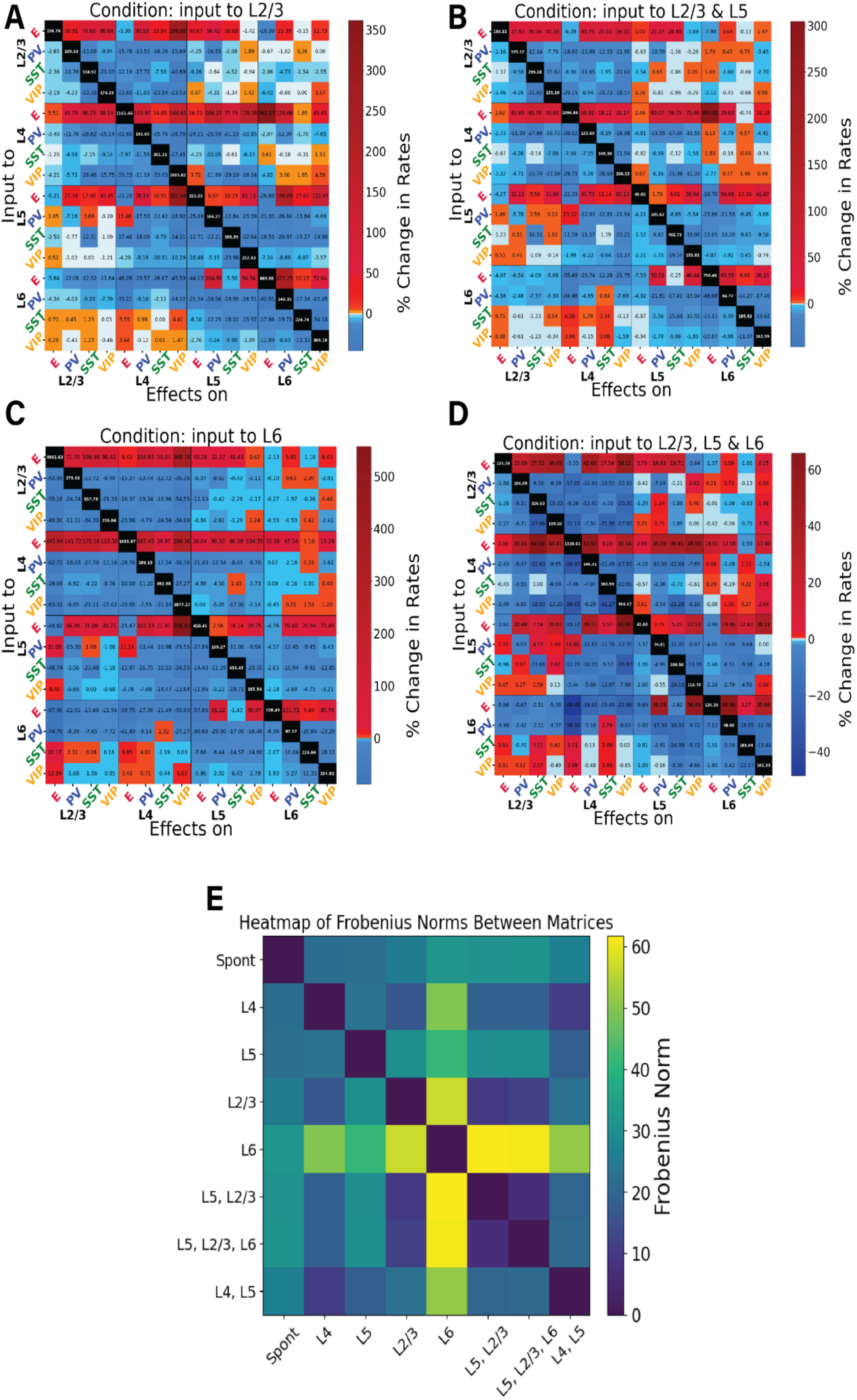
Perturbation analysis in four different feedback-driven states. Panels (A), (B), (C), (D) provide additional information on the data presented in Fig. 5. (A)-(D) Perturbation Input-output matrices for four distinct feedback states: A) Input to subset of E and PV neurons in layer 2/3. B) Input to subset of E and PV neurons in layer 2/3 and layer 5. C) Input to subset of E and PV neurons in layer 6. D) Input to subset of E and PV neurons in layer 2/3, 5 and 6. Within each condition, we delivered a 30 pA DC current to one population (Y-axis) and observed the resultant effects on the others (X-axis). We replicated this procedure for all 16 populations to derive each matrix. The matrices show the exact percentage changes of the firing rates compared to the situation pre-perturbation injection. (E) To quantify the differences between the conditions we used the Frobenius norm: we compute the pairwise distances between all the matrices to see which conditions are closer to each other.

**Figure S12:**
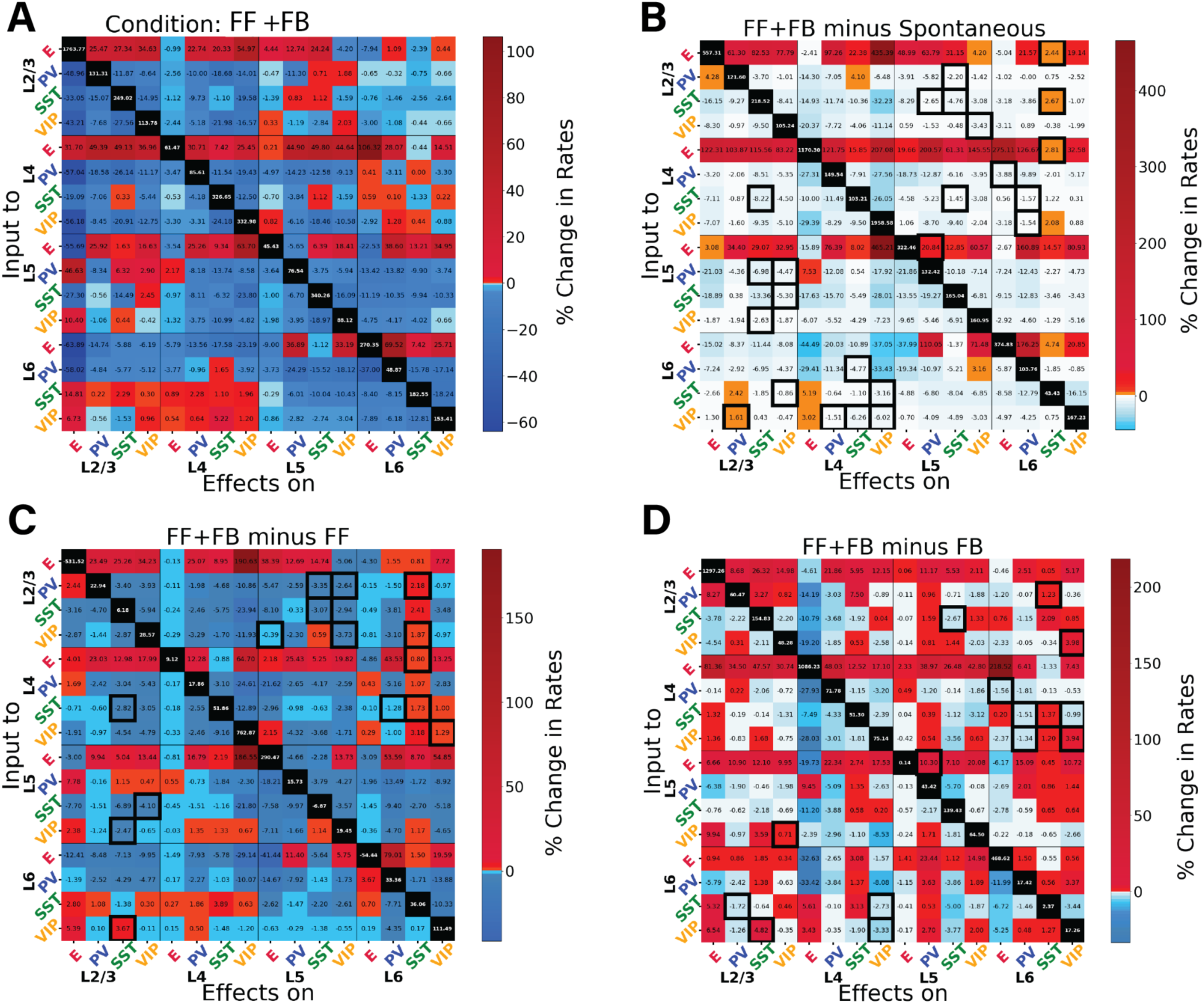
Perturbation analysis for the network in the feedback and feedforward inputs combined. Panels (A), (B), (C), (D) provide additional information on the data presented in Fig. 6. (A) Matrix of input-output relationships of the network. When both inputs to L4 and L5 are present, we delivered a 30 pA DC current to one population (Y-axis) and observed the resultant effects on the others (X-axis). We repeated this procedure for the 16 populations to obtain the matrix. The matrices show the exact percentage changes of the firing rates compared to the situation pre-perturbation injection. (B) Matrix showing the difference between the feedback & feedforward situation and the spontaneous condition. (C) Matrix showing the difference between the feedback & feedforward situation and the feedforward only condition. (D) Matrix showing the difference between the feedback & feedforward situation and the feedback only condition. The black contour square indicates when a change in sign between the two matrices occurred.

**Figure S13:**
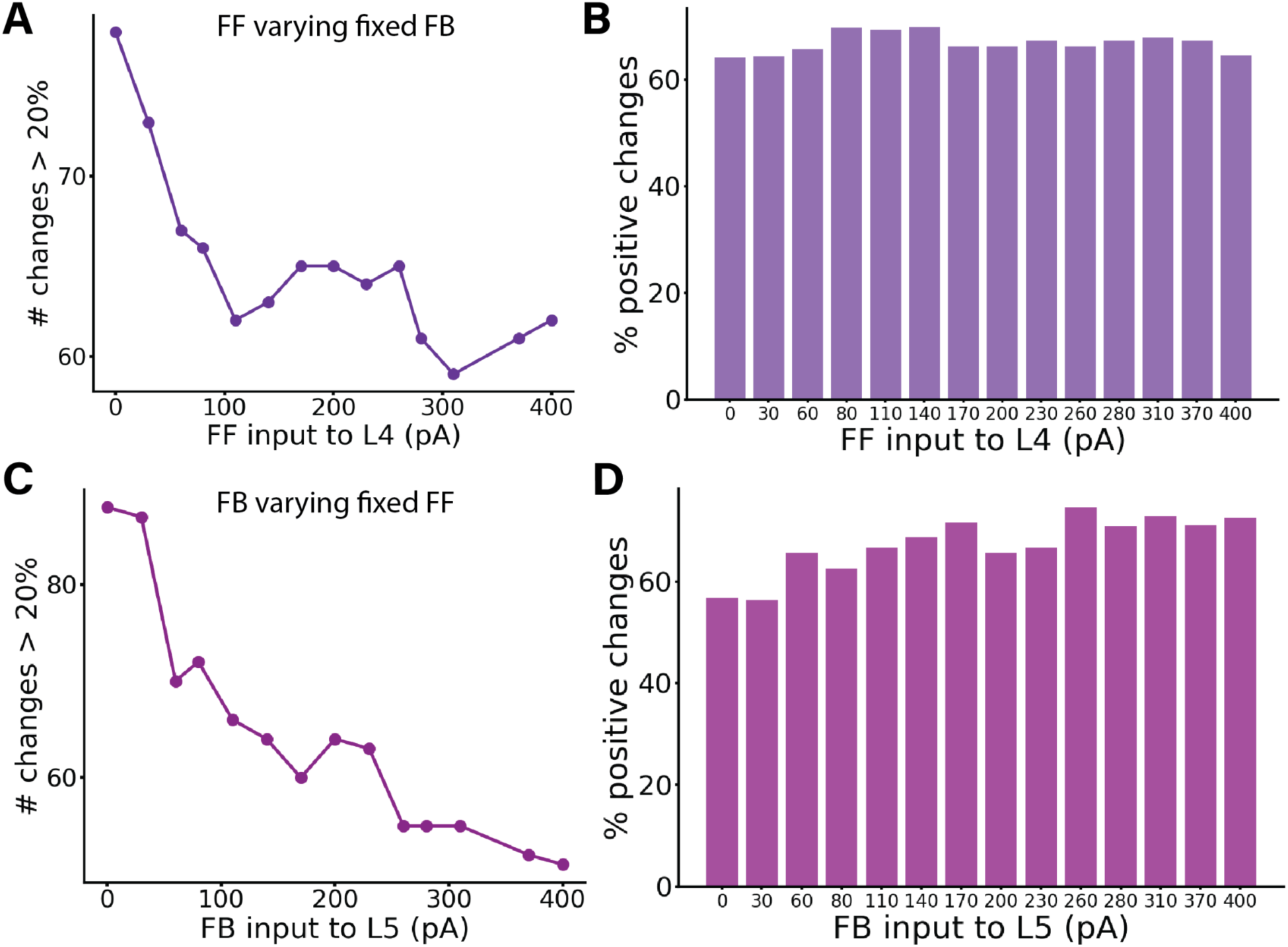
Perturbation Analysis in Different Conditions: A-B) Varying FF input while keeping FB constant. C-D) Varying FB input while keeping FB constant. (A) displays the number of changes >20% (or <-20%) in the corresponding perturbation matrix. Perturbation analysis was conducted for 14 different network conditions, defined by varying feedforward (FF) input to layer 4 while keeping feedback (FB) input to a subset of pyramidal cells and PV cells in layer 5 constant at 150 pA, resulting in a 16×16 matrix for each cell group, although these matrices are not displayed here. Each condition varied the input strength to neurons in layer 4 (25% E4 and 5% PV4), with values on the X-axis ranging from 0 to 400 pA. The Y-axis represents the number of alterations >20% (or <-20%) in the respective matrix (the sum of red and blue squares). Increased input to layer 4 results in fewer perturbation-induced changes in the firing rates of other populations. (B) presents percentages of positive changes in firing rates elicited by all possible perturbations in each network state, computed from the total number of changes shown in A. The complementary percentages in the panel also provide data for the negative changes. (C) illustrates the number of changes >20% (or <-20%) in the corresponding perturbation matrix. Perturbation analysis was conducted for 14 different network conditions, defined by varying feedback (FB) input to layer 5, and resulting in a 16×16 matrix for each cell group, though these matrices are not displayed here. Each condition varied the input strength to a subset of excitatory and PV neurons in layer 5, with values on the X-axis ranging from 0 to 400 pA. The Y-axis represents the number of t alterations >20% (or <-20%) in the respective matrix (the sum of red and blue squares). Increased input to layer 5 results in fewer perturbation-induced changes in the firing rates of other populations. (D) shows percentages of positive changes in firing rates elicited by all possible perturbations in each network state. The percentage is computed from the total number of changes presented in C. The complementary percentages for the panel also provide data for the negative changes.

**Figure S14:**
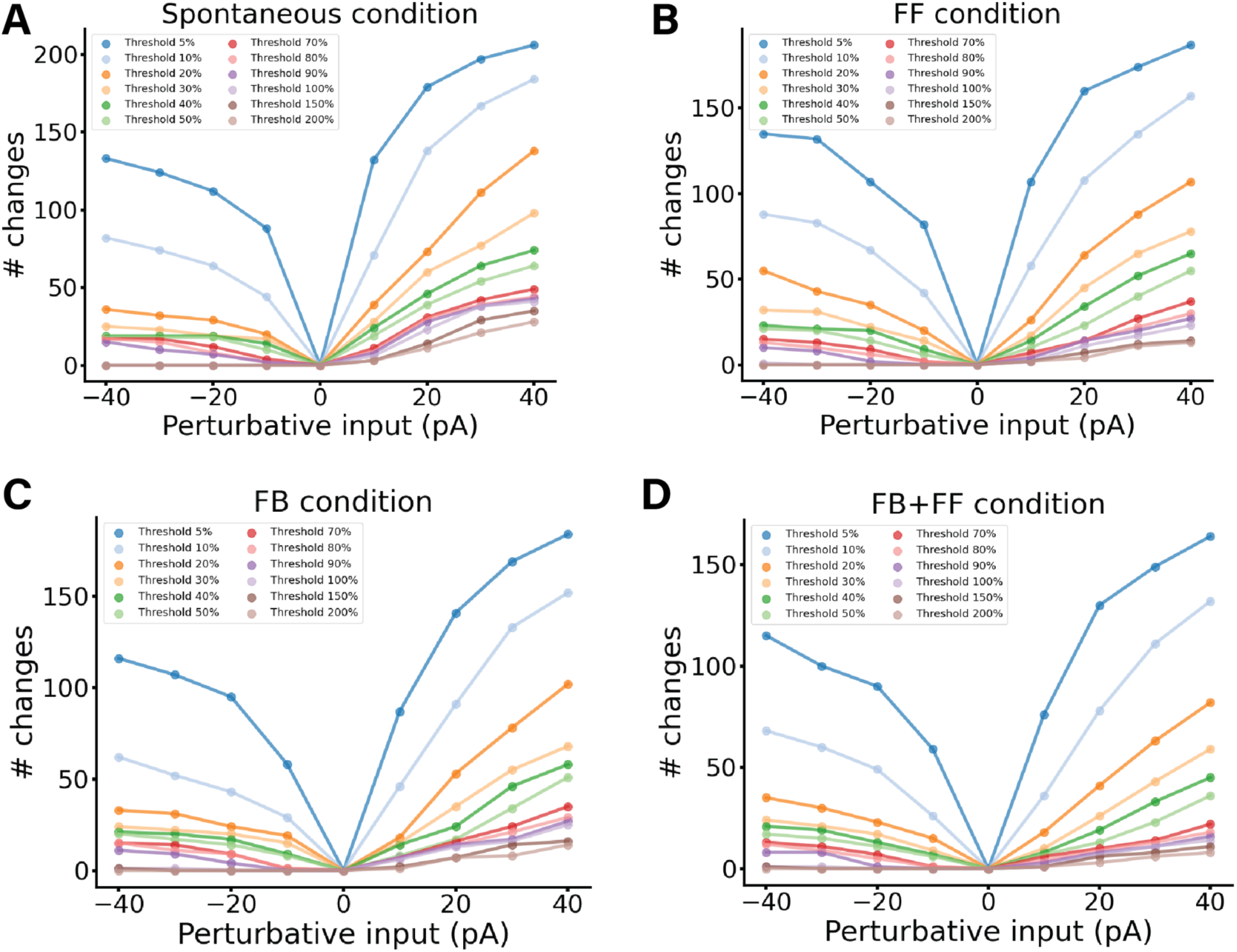
Perturbation Analysis in Different Conditions for different perturbation values. A) Spontaneous condition. The perturbation values used are [-40, −30, −20, −10, 10, 20, 30, 40] pA, for each value a perturbation analysis is carried out and a perturbation matrix is created. As usual to construct the matrix we delivered perturbative input to one population and observed its effect on others, repeating this for all 16 populations. We stimulated each subpopulation with a DC current of a given strength and monitored the resultant firing rate changes in all other subpopulations. Then for each matrix we counted the number of cells which changed their firing rate by more than the chosen threshold. Therefore we were able to draw the coloured lines in the plot. The threshold chosen for the figure are: 5%, 10%, 20%, 30%, 40%, 50%, 70%,80%, 90%, 100%, 150%, 200% indicating an increase or decrease of the firing rate of that quantity. B) Same analysis for Feedforward-driven state, wherein input is provided to a subset of L4 pyramidal cells and PV cells. C) Same analysis for Feedback-driven state, wherein input is provided to a subset of L5 pyramidal cells and PV cells. D) Same analysis for Feedforward and Feedback-driven state, wherein input is provided to a subset of L5 and L4 pyramidal cells and PV cells. For all condition the more perturbative input is injected the more the cells are changing their firing rate.

**Figure S15:**
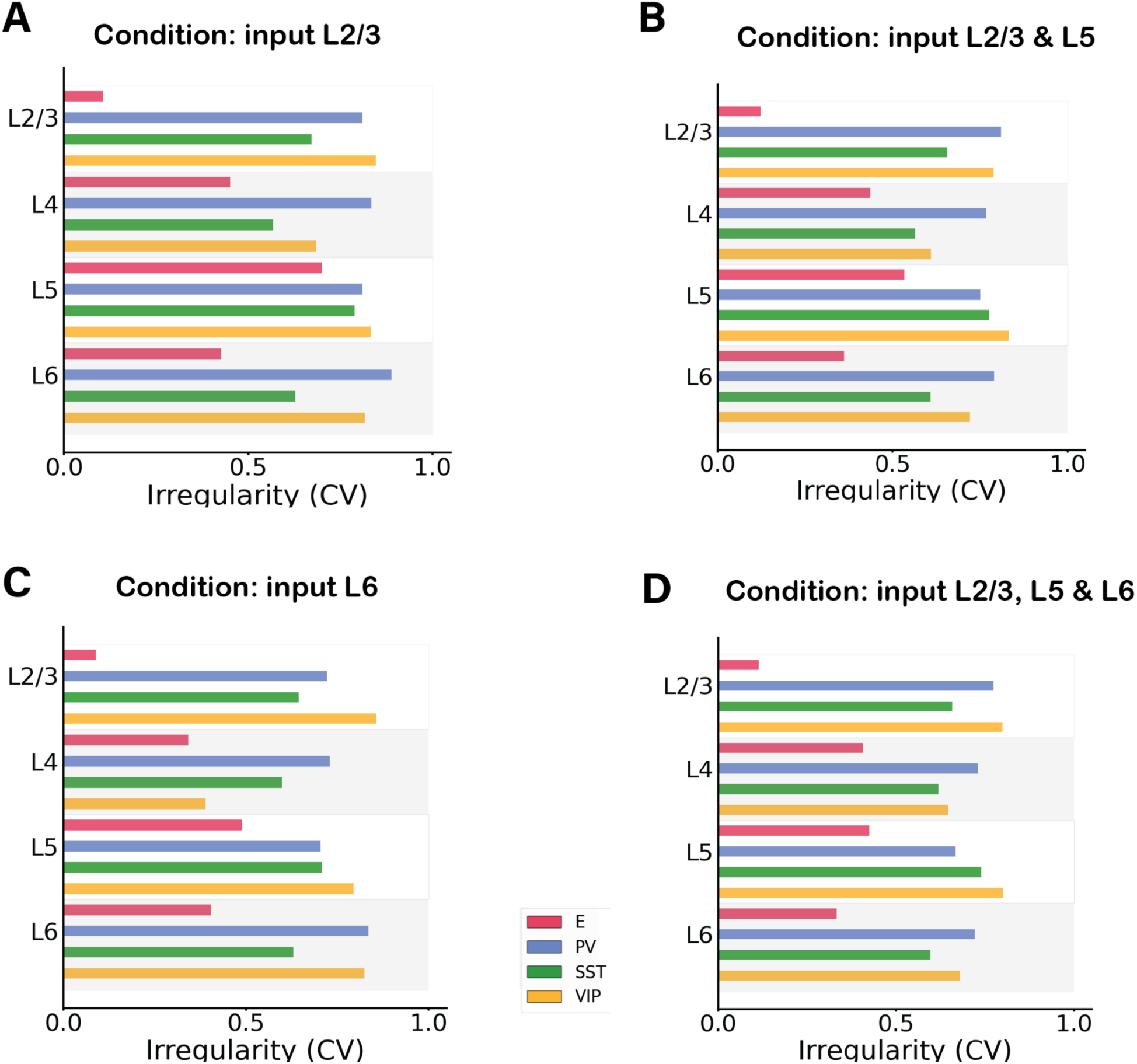
Irregularity of single-unit spike trains quantified by the coefficient of variation of the inter-spike intervals in four different feedback network condition. A) Condition: Feedback input to subset of E and PV neurons in layer 2/3. B) Condition: Input to subset of E and PV neurons in layer 2/3 and layer 5. C) Condition: Input to subset of E and PV neurons in layer 6. D) Condition: Input to subset of E and PV excitatory neurons in layer 2/3, 5 and 6. In all conditions most cells have a CV >0.5 or around 0.5 showing no synchrony. Excitatory cells in layer 2/3 have a low CV, caused by the firing rate activity being very low for those cells, and therefore making the CV harder to properly evaluate.

**Figure S16:**
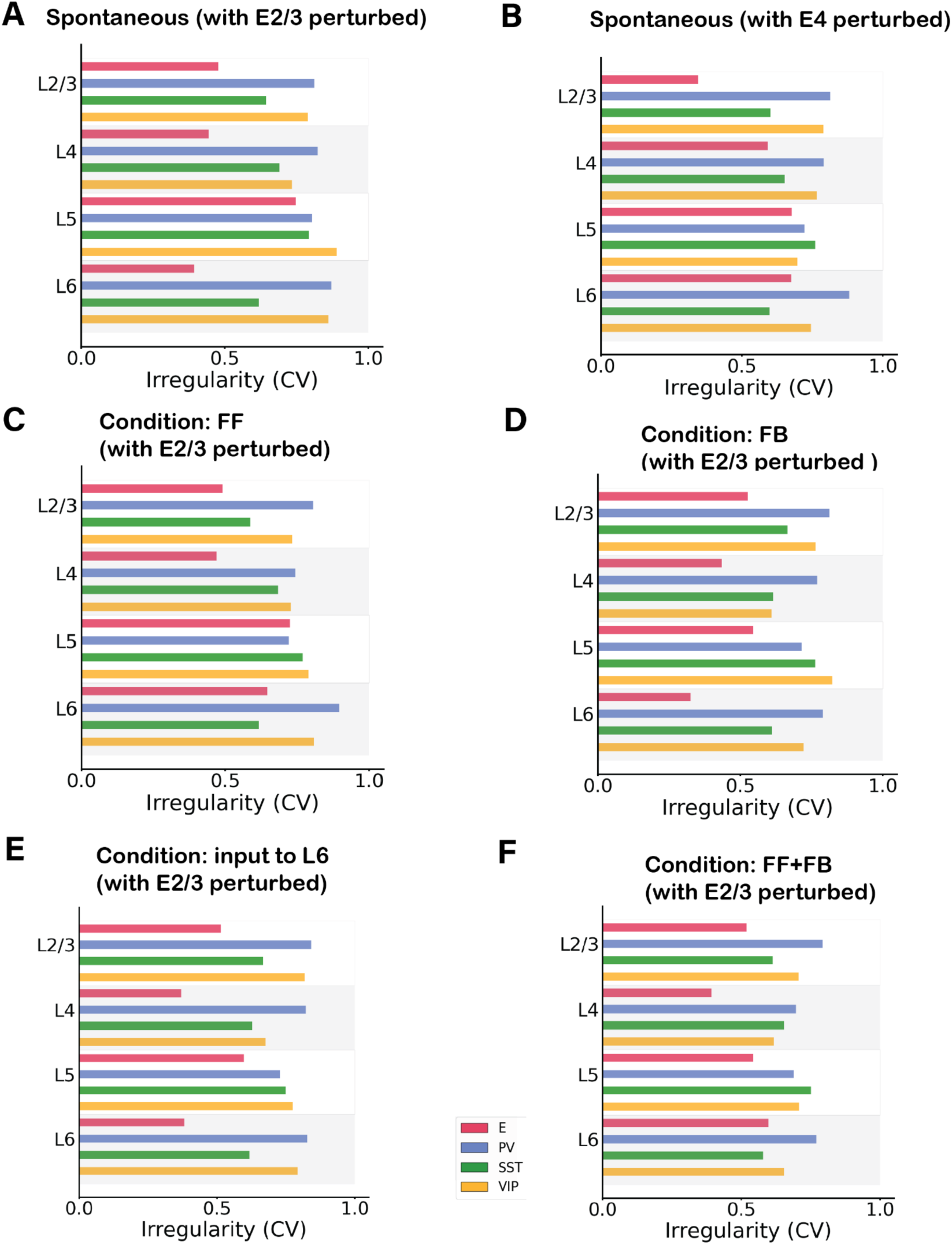
Irregularity of single-unit spike trains quantified by the coefficient of variation of the inter-spike intervals in 6 different network condition with a perturbative input. A) Condition: Spontaneous with perturbation of 30pA in excitatory neurons in layer 2/3. B) Condition: Spontaneous with perturbation of 30pA in excitatory neurons in layer 4. C) Condition: Feedforward input of 150pA injected in 25% excitatory cells and 5% PV cells in layer 4 and perturbative input to excitatory cells in layer 2/3. D) Condition: Feedback input of 150pA injected in 25% excitatory cells and 5% PV cells in layer 5 and perturbative input to excitatory cells in layer 2/3. E) Feedback input of 150pA injected in 25% excitatory cells and 5% PV cells in layer 6 and perturbative input to excitatory cells in layer 2/3. F) Condition: Feedforward and Feedback input combined, 150pA injected in 25% excitatory cells and 5% PV cells in layer 4 and 5 and perturbative input to excitatory cells in layer 2/3. In all conditions most cells have a CV >0.5 or around 0.5 showing no synchrony.

## Supplementary Tables

**Table 1:** Background noise rate that each group receives. Values represent the Poisson generator rates υ_bg_.

| Rate of Poisson generator $u_{bg}$ (Hz) | E | PV | SST | VIP |
| --- | --- | --- | --- | --- |
| L1 |  |  |  | 650 |
| L2/3 | 930 | 1460 | 870 | 1405 |
| L4 | 890 | 1984 | 2105 | 240 |
| L5 | 4740 | 930 | 530 | 870 |
| L6 | 1770 | 1170 | 885 | 1620 |

**Table 2:** Number of neurons in each layer. Ntot is the total number of cells in the column and can be defined arbitrarily for a simulation. The number of cells in each layer will scale accordingly [23]. We used Ntot=5000 for simulations.

|  | Number of neurons |
| --- | --- |
| L1 | $0.0192574218 * N_{tot}$ |
| L2/3 | $0.291088453 * N_{tot}$ |
| L4 | $0.237625904 * N_{tot}$ |
| L5 | $0.17425693 * N_{tot}$ |
| L6 | $0.297031276 * N_{tot}$ |

**Table 3:** Percentage of inhibitory cell types as a fraction of the total number of inhibitory cells in each layer. In each layer, the inhibitory cells represent 15% of the total number of neurons for that layer [23].

| Percentage (%) | PV | SST | VIP |
| --- | --- | --- | --- |
| L2/3 | 0.295918 | 0.214286 | 0.489796 |
| L4 | 0.552381 | 0.295238 | 0.152381 |
| L5 | 0.485714 | 0.428571 | 0.085714 |
| L6 | 0.458333 | 0.458333 | 0.083333 |

**Table 4:** Cell counts in the network for Ntot = 5000.

| Number of neurons | E | PV | SST | VIP |
| --- | --- | --- | --- | --- |
| L1 |  |  |  | 96 |
| L2/3 | 1236 | 65 | 47 | 107 |
| L4 | 1010 | 98 | 53 | 27 |
| L5 | 741 | 63 | 56 | 11 |
| L6 | 1263 | 102 | 102 | 19 |

**Table 5:** Membrane capacitance for each group of cells. The values are taken from the Allen database [23]. The same values are used for all simulations.

| $C_m$ (pF) | $E$ | $PV$ | $SST$ | $VIP$ |
| --- | --- | --- | --- | --- |
| $L1$ | | | | 37.11 |
| $L2/3$ | 123.41 | 70.95 | 82.34 | 41.23 |
| $L4$ | 80.16 | 81.21 | 132.86 | 40.3 |
| $L5$ | 149.43 | 70.9 | 52.32 | 59.29 |
| $L6$ | 99.96 | 49.65 | 96.09 | 65.87 |

**Table 6:** Leak conductance for each group of cells [23].

| $g_L$ (nS) | $E$ | $PV$ | $SST$ | $VIP$ |
| --- | --- | --- | --- | --- |
| $L1$ | | | | 4.07 |
| $L2/3$ | 2.47 | 9.49 | 3.17 | 6.4 |
| $L4$ | 5.16 | 9.19 | 7.96 | 1.87 |
| $L5$ | 16.66 | 5.21 | 3.43 | 6.52 |
| $L6$ | 5.88 | 6.86 | 2.99 | 6.09 |

**Table 7:**
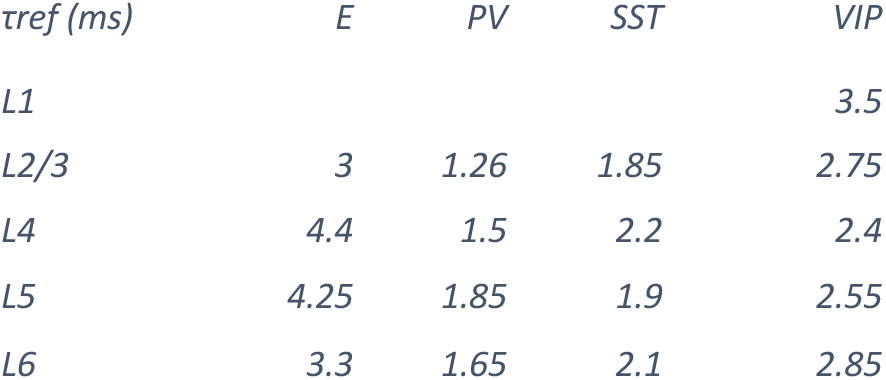
Refractory period for each group of cells [23].

| $\tau_{ref}$ (ms) | $E$ | $PV$ | $SST$ | $VIP$ |
| --- | --- | --- | --- | --- |
| $L1$ | | | | 3.5 |
| $L2/3$ | 3 | 1.26 | 1.85 | 2.75 |
| $L4$ | 4.4 | 1.5 | 2.2 | 2.4 |
| $L5$ | 4.25 | 1.85 | 1.9 | 2.55 |
| $L6$ | 3.3 | 1.65 | 2.1 | 2.85 |

**Table 8:** Resting membrane potential for each group of cells [23].

| $V_{rest}$ (mV) | $E$ | $PV$ | $SST$ | $VIP$ |
| --- | --- | --- | --- | --- |
| $L1$ | | | | -65.5 |
| $L2/3$ | -80.97 | -82.35 | -69.16 | -67.94 |
| $L4$ | -72.53 | -70.45 | -74.2 | -63.14 |
| $L5$ | -68.28 | -77.5 | -70.01 | -72.00 |
| $L6$ | -77.5 | -76.42 | -62.99 | -78.85 |

**Table 9:** Spike threshold for each group of cells [23].

| *V<sub>th</sub>* (mV) | *E* | *PV* | *SST* | *VIP* |
| --- | --- | --- | --- | --- |
| *L1* |  |  |  | -40.20 |
| *L2/3* | -40.53 | -56.32 | -39.95 | -41.34 |
| *L4* | -47.63 | -44.23 | -44.07 | -40.89 |
| *L5* | -40.55 | -51.2 | -47.38 | -51.2 |
| *L6* | -42.31 | -49.06 | -37.19 | -44.81 |

**Table 10:**
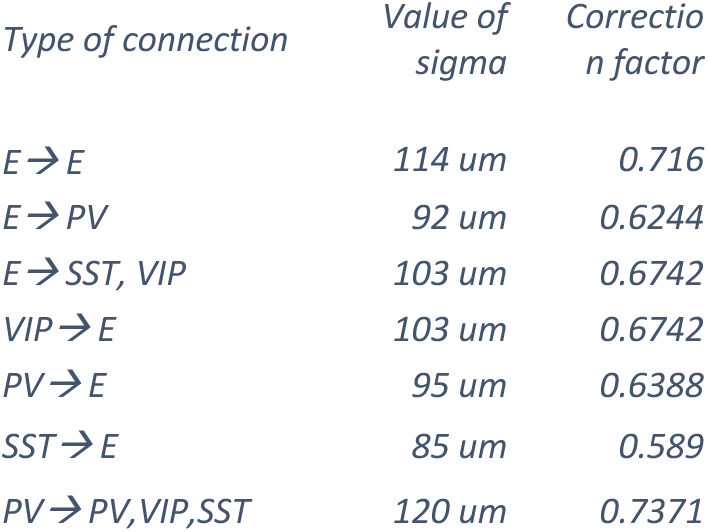
Corrections of connection probability as a function of the Gaussian width for each connection type.

